# MAIT cell responses to intracellular and extracellular pathogens are mediated by distinct antigen presenting cells

**DOI:** 10.1101/2025.05.03.652000

**Authors:** Dominic Haas, Jonathan Melamed, Gabrielle LeBlanc, Nikhita Arun, Andrew J. Perkowski, Jeffrey Aubé, Matthias Mack, Michael G. Constantinides

## Abstract

Mucosal-associated invariant T (MAIT) cells recognize microbial derivatives of riboflavin synthesis presented by the MHC class I-related (MR1) molecule. Although these metabolites are highly conserved among bacteria, the cells that present them remain unknown. Here, we show type-17 MAIT cells respond to diverse isolates of the extracellular pathogen *Acinetobacter baumannii* and promote bacterial clearance. Both hematopoietic and non-hematopoietic cells mediate MR1 presentation within the lungs and mediastinal lymph nodes (meLNs). Conversely, the type-1 MAIT cell response to the intracellular pathogen *Francisella tularensis* requires MR1 presentation by type-2 conventional dendritic cells (cDC2s) within meLNs and ablation of these cells or their expression of MR1 renders animals more susceptible to the infection. Although MR1 is broadly expressed at homeostasis, *A. baumannii* enhances MR1 on macrophages and fibroblasts, while *F. tularensis* increases expression on cDC2s. These results demonstrate that microbial tropism dictates which APCs mediate MR1 presentation of metabolites, revealing alternative therapeutic approaches.

## Introduction

Adaptive immunity is initiated by processing microbial proteins into peptides that are presented on major histocompatibility complex (MHC) molecules.^1^ MHC class I (MHC-I) presents endogenous peptides on all nucleated cells and facilitates cross-presentation of exogenous peptides primarily by dendritic cells (DCs), both of which result in the activation of cytotoxic CD8^+^ T cells.^2^ MHC-II stimulates CD4^+^ T cells through the presentation of peptides derived from extracellular and intravesicular pathogens on antigen-presenting cells (APCs), including DCs, macrophages, and B cells.^3^ In addition to the peptides presented by polymorphic MHC proteins, non-peptidic antigens are presented by monomorphic MHC-Ib molecules to unconventional T cells.^4, 5^ The cluster of differentiation 1 (CD1) family member CD1d presents lipids to invariant natural killer T (iNKT) cells, while the MHC-I related (MR1) molecule presents metabolites to mucosal-associated invariant T (MAIT) cells.^4, 5^ Although the phylogenetic conservation of CD1d and MR1 suggests that presentation of non-peptidic microbial antigens has been necessary throughout mammalian evolution,^6^ the APCs that mediate this remain unclear.

MAIT cells express semi-invariant T cell receptors (TCRs; Vα19-Jα33 in mice and Vα7.2-Jα33/20/12 in humans) that recognize microbial derivatives of riboflavin (vitamin B2) synthesis presented by MR1.^7, 8^ While animals acquire riboflavin from their diets, most bacteria and fungi synthesize this vitamin to generate the coenzymes flavin adenine dinucleotide and flavin mononucleotide.^9, 10^ Consequently, MAIT cells produce interferon gamma (IFN-γ), interleukin-17A (IL-17A), and/or tumor necrosis factor (TNF) in response to many respiratory pathogens, including *Aspergillus* species, *Haemophilus influenzae, Mycobacterium tuberculosis*, and *Streptococcus pneumoniae*, with protection in murine models demonstrated for *Francisella tularensis, Klebsiella pneumoniae*, and *Legionella longbeachae*.^11–20^ MAIT cells are the most abundant unconventional T cells in most human tissues,^5, 8^ representing up to 8% of pulmonary T cells in healthy individuals,^14^ although other MR1-restricted T cells with diverse TCRs are also present in humans and mice.^21–23^ A single-nucleotide polymorphism in the *Mr1* gene is associated with increased susceptibility to tuberculosis and an individual with a nonfunctional MR1 variant experienced recurrent bacterial and viral infections, demonstrating that MR1-resticted T cells contribute to human health.^24, 25^

MR1 is the only MHC protein that presents metabolites, displaying molecules from both intracellular and extracellular microbes.^26, 27^ At steady state, MR1 predominantly resides within the endoplasmic reticulum (ER)-Golgi compartment in a ligand-receptive conformation that is stabilized by chaperone proteins.^26, 27^ During infection, microbial metabolites are loaded within the antigen binding site of MR1, inducing a conformational change that releases chaperone proteins and enables trafficking to the plasma membrane.^26, 27^ Self or environmental antigens are also thought to facilitate a low level of constitutive MR1 surface expression and can be exchanged for microbial metabolites within endosomes or phagosomes following internalization of MR1.^26–28^ The ubiquitous expression of MR1 and its retention within the ER-Golgi compartment has made identification of MR1-presenting cells challenging.

Although both intracellular and extracellular pathogens synthesize riboflavin, greater MAIT cell responses have been reported to intracellular bacteria, including *F. tularensis, L. longbeachae,* and *Salmonella enterica* serovar Typhimurium, which expand MAIT cells by more than 100-fold.^18–20, 29^ Conversely, the extracellular bacteria *K. pneumoniae*, *S. pneumoniae,* and *Staphylococcus epidermidis* increase MAIT cell abundance by less than 5-fold.^10, 30, 31^ Since the most potent MAIT cell agonist, the riboflavin derivative 5-(2-oxopropylideneamino)-6-D-ribitylaminouracil (5-OP-RU), degrades within 30 minutes unless stabilized by binding to MR1,^32^ intracellular microbial replication may facilitate MR1 loading. Therefore, we sought to establish whether an extracellular pathogen could elicit a strong MAIT cell response dependent on MR1 presentation.

Here, we determine that type-17 MAIT (MAIT17) cells respond robustly to diverse strains of the extracellular bacterial pathogen *Acinetobacter baumannii*, which causes nosocomial infections and is classified as an urgent threat by the U.S. Centers for Disease Control and Prevention due to extensive antibiotic resistance.^33^ During *A. baumannii* pneumonia, MAIT17 cells are the primary IL-17A producers and promote bacterial clearance. MR1 presentation is necessary for the MAIT cell response to both *A. baumannii* and the intracellular bacterial pathogen *F. tularensis,* permitting comparison of metabolites sourced from distinct locations. Using a novel *Mr1^FLAG^* reporter strain, we establish that MR1 protein is most highly expressed by alveolar macrophages (AMΦs) and non-hematopoietic cells within the lungs at steady state. Their expression of MR1 increases during infection with *A. baumannii*, but not *F. tularensis*, which instead enhances MR1 expression by type-2 conventional dendritic cells (cDC2s). While hematopoietic and non-hematopoietic cells mediate presentation of *A. baumannii* metabolites, the type-1 MAIT (MAIT1) cell response to *F. tularensis* requires MR1 presentation by cDC2s and ablation of these cells or their expression of MR1 renders animals more susceptible to the infection. These results demonstrate that microbial tropism dictates which APCs mediate MR1 presentation of metabolites.

## Results

### MAIT17 cells provide immunity to *A. baumannii*

While the innate immune response to pulmonary *A. baumannii* infection has been characterized, the adaptive immune response remains undescribed.^34–36^ To interrogate a potential role for T lymphocytes, we inoculated wild-type (WT) C57BL/6J mice with 10^8^ colony-forming units (CFUs) of the human pathogenic isolate *A. baumannii* strain 17978 retropharyngeally (RPh). Flow cytometry analysis of the lungs 1 week later revealed that MAIT cells expanded the most among T lymphocytes, increasing ∼70-fold in number compared to uninfected animals **(Fig. 1A & S1A)**. By comparison, CD4^+^ and CD8^+^ conventional T cells and other innate-like T cells (γδ and iNKT) expanded less than 5-fold. Administration of decreasing inoculums indicated that the MAIT cell response to *A. baumannii* was dose-dependent, with the number of MAIT cells roughly proportional to bacterial CFUs **(Fig 1B)**. Significant accumulation of MAIT cells occurred 5 days after *A. baumannii* inoculation **(Fig. 1C)**, which is comparable to the MAIT cell expansion following pulmonary *F. tularensis* infection.^18, 19^ MAIT cells that responded to *A. baumannii* expressed the transcription factor RORγt and the costimulatory molecule ICOS, which characterize MAIT17 cells, but not the MAIT1-associated chemokine receptor CXCR3 **(Fig. 1D-F)**.^37^ 7 days post-inoculation, MAIT17 cells were the dominant IL-17A-producing T cells within the lungs **(Fig. 1G-H)**, emphasizing their contribution during *A. baumannii* infection.

**Figure 1.**
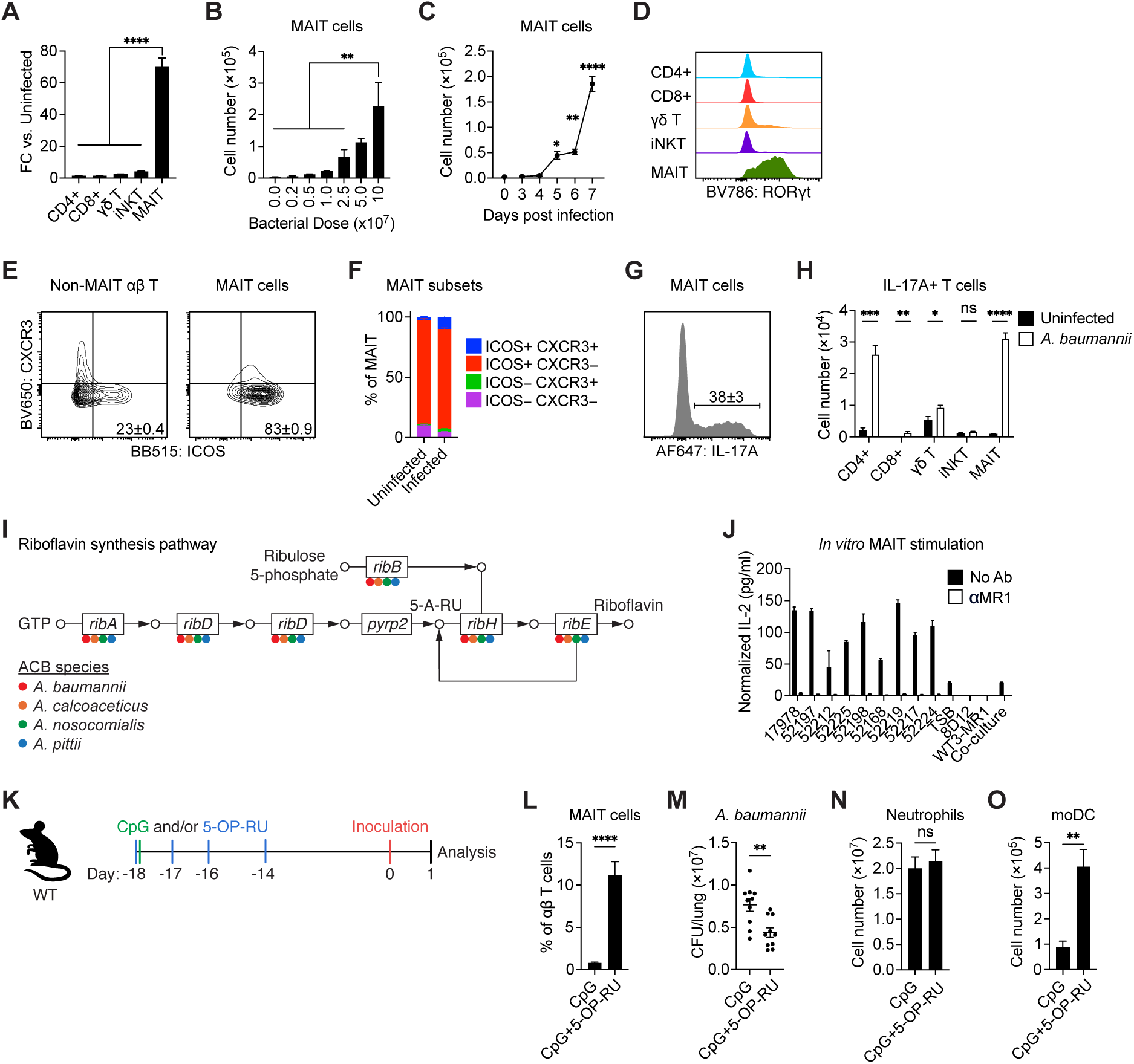
MAIT17 cells provide immunity to *A. baumannii*. (A) Fold change in various T lymphocyte subsets 7 DPI *A. baumannii* compared to average counts in uninfected mice (n=4-5/group). (B) Number of MAIT cells following various RPh doses of *A. baumannii* (n=3-6/group). (C) Number of MAIT cells 3-7 DPI *A. baumannii* (n=4-5/group). Asterisks represents significance to D0 MAIT counts. (D) Representative flow staining of RORγt on various lung T lymphocytes 7 DPI *A. baumannii.* (E and F) Representative (E) and summary (F) flow staining of CXCR3 and ICOS on MAIT cells and all other αβ T cells for uninfected or mice 7 DPI *A. baumannii* (n=5-9/group). Data represents average ± SEM. (G) Representative flow staining of IL-17A on MAIT cells of mice 7 DPI *A. baumannii* following *ex vivo* PMA/ionomycin re-stimulation (n=9/group). Data represents average ± SEM. (H) Number of IL-17A positive cells across various lymphocyte subsets following *ex vivo* PMA/ionomycin re-stimulation for uninfected or mice 7 DPI *A. baumannii* (n= 5-9/group). (I) Schematic representing presence of riboflavin synthesis genes in species of *Acinetobacter* denoted with the appropriate color. Dot underneath gene represents >90% conservation across strains of the ACB species. (J) Supernatant IL-2 following 20-hour co-culture of MR1-overexpressing cell line (WT3-MR1) with murine MAIT hybridoma (8D12) with or without blocking MR1 antibody. Co-culture spiked with 1:100 dilution of various *A. baumannii* strain supernatants or media control (TSB: tryptic soy broth). IL-2 levels normalized to 10^9^ CFU. (K) Schematic for boosting MAIT cells in the lung. On -18 DPI, 20μg CpG and 50nmol 5-OP-RU are co-administered followed by 5-OP-RU only on -17, -16, and -14 DPI. (L-O) Percentage of MAIT cells (L), CFU per lung (M), absolute neutrophil count (N), and absolute moDC count (O) 1 DPI with 1×10^8^ CFU *A. baumannii* compared between mice treated with CpG and CpG+5-OP-RU (n=10 mice/group). Graphs indicate means ±SEM. One-way ANOVA (A, B, C), Student’s *t*-tests (H, L, M), and Mann Whitney U tests (N, O) were performed. Statistics: ns, not significant (*P* > 0.05); \**P* < 0.05; \*\**P* < 0.01; \*\*\**P* < 0.001, \*\*\*\**P* < 0.0001.

*A. baumannii* is a member of the *Acinetobacter calcoaceticus-baumannii* (ACB) complex that also includes *A. calcoaceticus*, *A. nosocomialis*, and *A. pittii*, the latter 2 of which are human pathogens.^38, 39^ The *rib* operon that encodes riboflavin synthesis enzymes was highly conserved among these *Acinetobacter* species, with the associated genes present in greater than 90% of isolates **(Fig. 1I)**. Absence of the *pyrp2* gene suggests these bacteria synthesize riboflavin from ribulose 5-phosphate rather than guanosine-5′-triphosphate (GTP) to generate the 5-OP-RU precursor 5-amino-6-D-ribitylaminouracil (5-A-RU). We determined whether MAIT cells respond to other isolates of *A. baumannii* by adding supernatants to co-cultures of WT3 murine fibroblasts overexpressing murine MR1 (WT3-MR1) and 8D12 murine MAIT hybridoma cells,^40, 41^ which release IL-2 upon activation **(Fig. S1B)**. All *A. baumannii* isolates induced IL-2 production, which was abrogated by the addition of an MR1-blocking antibody **(Fig. 1J)**. Thus, MAIT cells recognize a conserved *A. baumannii* antigen that is presented on MR1.

Given the responsiveness of MAIT cells to *A. baumannii*, we sought to establish whether they contribute to immunity. Although the frequency of MAIT cells in human lungs is 4% of T lymphocytes on average and as high as 8% in some individuals,^14^ MAIT cells account for 0.5-1% of pulmonary T lymphocytes in C57BL/6J mice. To increase the abundance of murine MAIT cells, we administered the riboflavin derivative 5-OP-RU with or without the Toll-like receptor 9 agonist CpG oligodeoxynucleotides RPh **(Fig. 1K)**, which has previously been shown to selectively expand MAIT cells.^29^ Treatment with 5-OP-RU and CpG increased MAIT cell abundance 10-fold over CpG alone **(Fig. 1L)**, which reduced the pulmonary burden of *A. baumannii* 24 hours post-inoculation **(Fig. 1M)**. While neutrophils contribute to the clearance of *A. baumannii*,^42, 43^ their abundance was unchanged by the increase in MAIT cells **(Fig. 1N & S1C-D)**. However, treatment with 5-OP-RU and CpG increased monocyte-derived DCs (moDCs) **(Fig. 1O)**, which is consistent with the finding that MAIT cells promote differentiation of monocytes into moDCs through their production of granulocyte-macrophage colony-stimulating factor.^44^ Together, these results indicate that MAIT17 cells are the most responsive T lymphocytes during *A. baumannii* pneumonia and promote the clearance of this pathogen.

### Intracellular and extracellular pathogens induce distinct MAIT cell programs

Although the majority of *A. baumannii* strains are noninvasive, strain 17978 can invade lung epithelial cells but does not replicate intracellularly.^45^ While the type-17 response to this strain is consistent with an extracellular microbe, we compared the responding MAIT cells to those induced by the live vaccine strain (LVS) of the intracellular pathogen *F. tularensis*. 7 days following RPh inoculation with *A. baumannii* strain 17978 (10^8^ CFUs) or *F. tularensis* LVS (2×10^3^ CFUs), pulmonary MAIT cells were isolated by fluorescence-activated cell sorting (FACS) and transcriptionally profiled using single-cell RNA sequencing (scRNA-seq). *A. baumannii*-responsive MAIT cells predominantly grouped with MAIT cells from uninfected animals, which are primarily MAIT17 cells within the lungs,^19^ while *F. tularensis*-responsive MAIT cells clustered separately **(Fig. 2A)**. The MAIT17 and MAIT1 transcriptional signatures were highly enriched in cells that responded to *A. baumannii* and *F. tularensis*, respectively **(Fig. 2B)**. Modest enrichment of the MAIT17 signature in *F. tularensis*-responsive MAIT cells is consistent with prior studies that have identified both T-bet^+^ MAIT1 cells and RORγt^+^ T-bet^+^ MAIT1/17 cells during *F. tularensis* infection **(Fig. 2B)**.^19, 46^ Louvain clustering identified 11 transcriptional states, including clusters that were enriched in *A. baumannii*-responsive (clusters 0, 2, & 9) or *F. tularensis*-responsive (clusters 4, 5, 8, & 10) MAIT cells **(Fig 2C-D)**. Although *A. baumannii*-responsive clusters exclusively expressed the MAIT17 signature, the *F. tularensis*-responsive clusters 4, 5, and 8 expressed both MAIT1 and MAIT17 **(Fig. S2A)**. *F. tularensis*-responsive cells and their associated clusters also expressed transcriptional signatures associated with the S, G_2_, and M cell cycle stages **(Fig. 2E & S2A-B)**, which is consistent with an ongoing *F. tularensis* infection.^18^ *A. baumannii*-responsive MAIT cells expressed residency and tissue repair signatures, which concurs with the higher tissue residency index of MAIT17 cells and their association with wound healing **(Fig. 2E & S2A-B)**.^10, 47^ These cells also expressed the MAIT17-associated genes *Rorc*, *Il17a*, *Il1r1*, *Ccr6*, *Sdc1*, and *Il7r* **(Fig. 2F)**, while *F. tularensis*-responsive MAIT cells expressed the MAIT1-associated genes *Tbx21*, *Ifng*, *Gzmb*, *Il12rb2*, *Cxcr3*, and *Klrg1* **(Fig. 2G)**.^31, 37^ Therefore, the extracellular pathogen *A. baumannii* induces a MAIT17 response that is transcriptionally distinct from the predominantly MAIT1 response to the intracellular pathogen *F. tularensis*.

**Figure 2.**
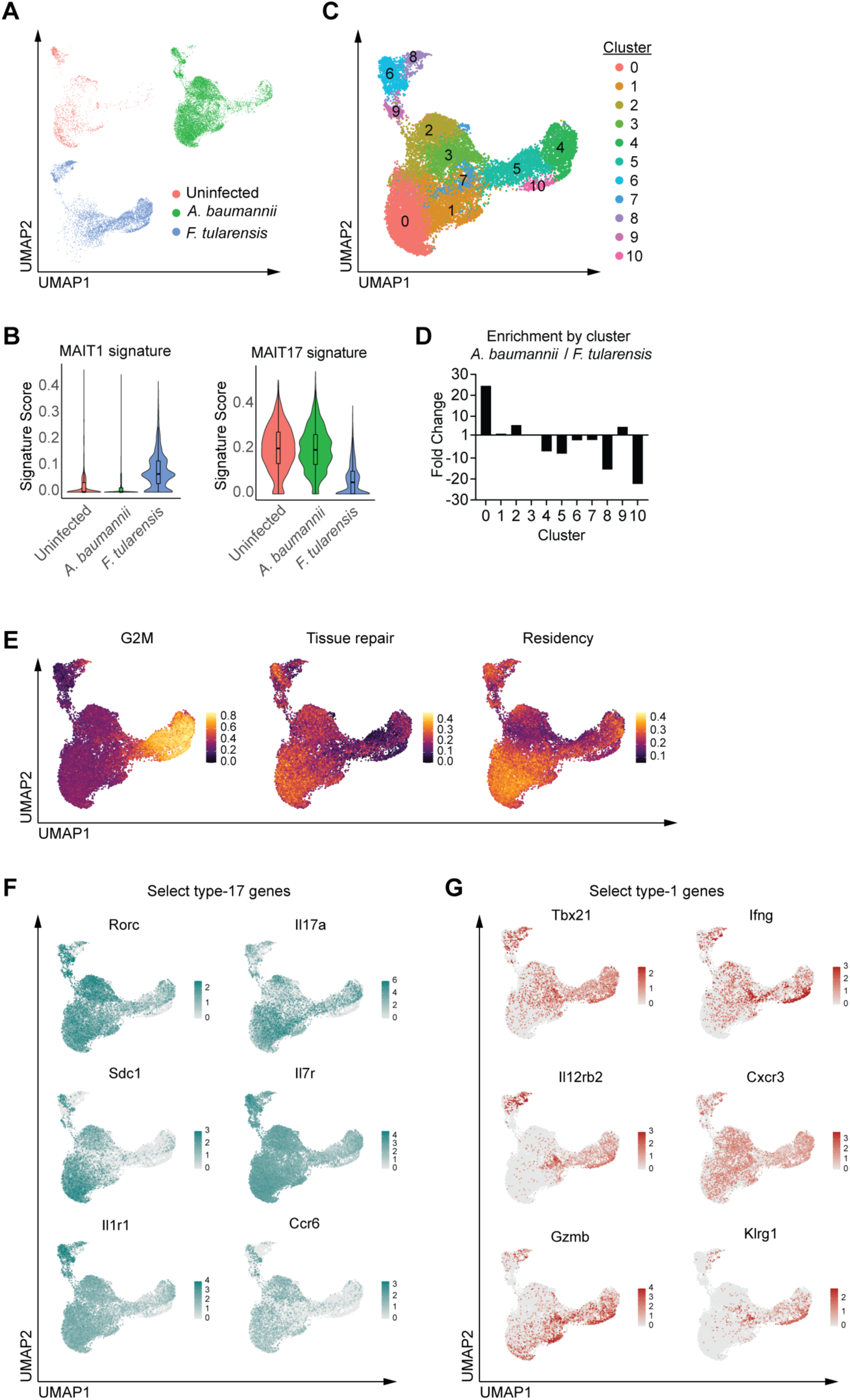
Intracellular and extracellular pathogens induce distinct MAIT cell programs. scRNA-seq data of MAIT cells sorted from uninfected (1482 cells), 7 DPI RPh *A. baumannii* (11690 cells), and 7 DPI RPh *F. tularensis* (5470 cells) using 10X Genomics platform. Plots were generated by combining scRNA-seq libraries from these three samples. (A) UMAP plot split by sample. (B) Enrichment for MAIT1 signature (left) MAIT17 signature (right) split by sample. (C) UMAP plot displaying the unbiased clustering of sequenced MAIT cells using Louvain algorithm. (D) Differential distribution of cells between *A. baumannii* and *F. tularensis* samples. A quotient of cells in each cluster over total cells per sample was calculated to determine fold change between samples. (E) UMAP plot displaying enrichment for G2M, tissue repair, and residency signatures. (F-G) UMAP plot displaying select type-17 genes (F) and select type-1 genes (G).

### MR1 protein is broadly expressed and increased during infection

While microbial metabolites facilitate trafficking of MR1 to the plasma membrane,^26, 27^ pathogens can also suppress MR1 presentation,^48^ so we sought to determine how *A. baumannii* and *F. tularensis* affect MR1 expression. Although multiple cell types are capable of MR1-mediated antigen presentation *in vitro*, including macrophages, DCs, monocytes, B cells, and airway epithelial cells,^12, 49^ the cells that express MR1 *in vivo* have not been identified because commercially available anti-MR1 antibodies poorly resolve murine MR1 and stain non-specifically during infection **(Fig. S3A)**. To quantify MR1 expression, we appended a glycine-serine-glycine linker, biotin acceptor peptide, and triple FLAG sequences to the final exon of *Mr1*, retaining the 3′ untranslated region that regulates messenger RNA stability **(Fig. 3A)**. Use of peptide tags preserves glycosylation of the MR1 protein and its post-transcriptional regulation by extracellular signal-regulated kinases.^50, 51^ We validated the resulting *Mr1^FLAG^* allele using CD4^+^ CD8^+^ thymocytes, which express high levels of MR1.^52^ Since the peptide tag prevented MR1 surface expression **(Fig. S3B)**, we stained anti-FLAG intracellularly, which resolved substantially more MR1-FLAG expression than anti-MR1 **(Fig. 3B)**. In the lungs of uninfected *Mr1^FLAG/+^* mice, AMΦs expressed the greatest amount of MR1-FLAG among hematopoietic cells **(Fig. 3C)**, which agrees with prior work that identified *Mr1* as one of the most highly expressed genes in macrophages relative to DCs.^53^ Infection with *A. baumannii* or *F. tularensis* modestly enhanced MR1-FLAG expression by most hematopoietic cells, with similar levels observed on B cells, monocytes, and monocyte-derived DCs (moDCs) following either pathogen **(Fig. 3C-M)**. However, *A. baumannii* increased MR1-FLAG expression by AMΦs and interstitial macrophages (IMΦs) more than *F. tularensis* **(Fig. 3F-I)**. Conversely, *F. tularensis* upregulated MR1-FLAG on cDCs more than *A. baumannii*, with the greatest expression observed by MAR1^+^ inflammatory cDC2s (inf-cDC2s) **(Fig. 3K-L)**.^54^ Non-hematopoietic cells within the lungs also expressed MR1-FLAG, with fibroblasts displaying the highest levels **(Fig. 3N-P)**.^55^ Inoculation with *A. baumannii*, but not *F. tularensis*, enhanced fibroblast expression of MR1-FLAG, with the largest increases by adventitial and CD249^+^ alveolar fibroblasts **(Fig. 3Q-T)**. These results demonstrate that MR1 is most highly expressed by AMΦs and non-hematopoietic cells at homeostasis and infections with *A. baumannii* and *F. tularensis* increase MR1 expression on different cells.

**Figure 3.**
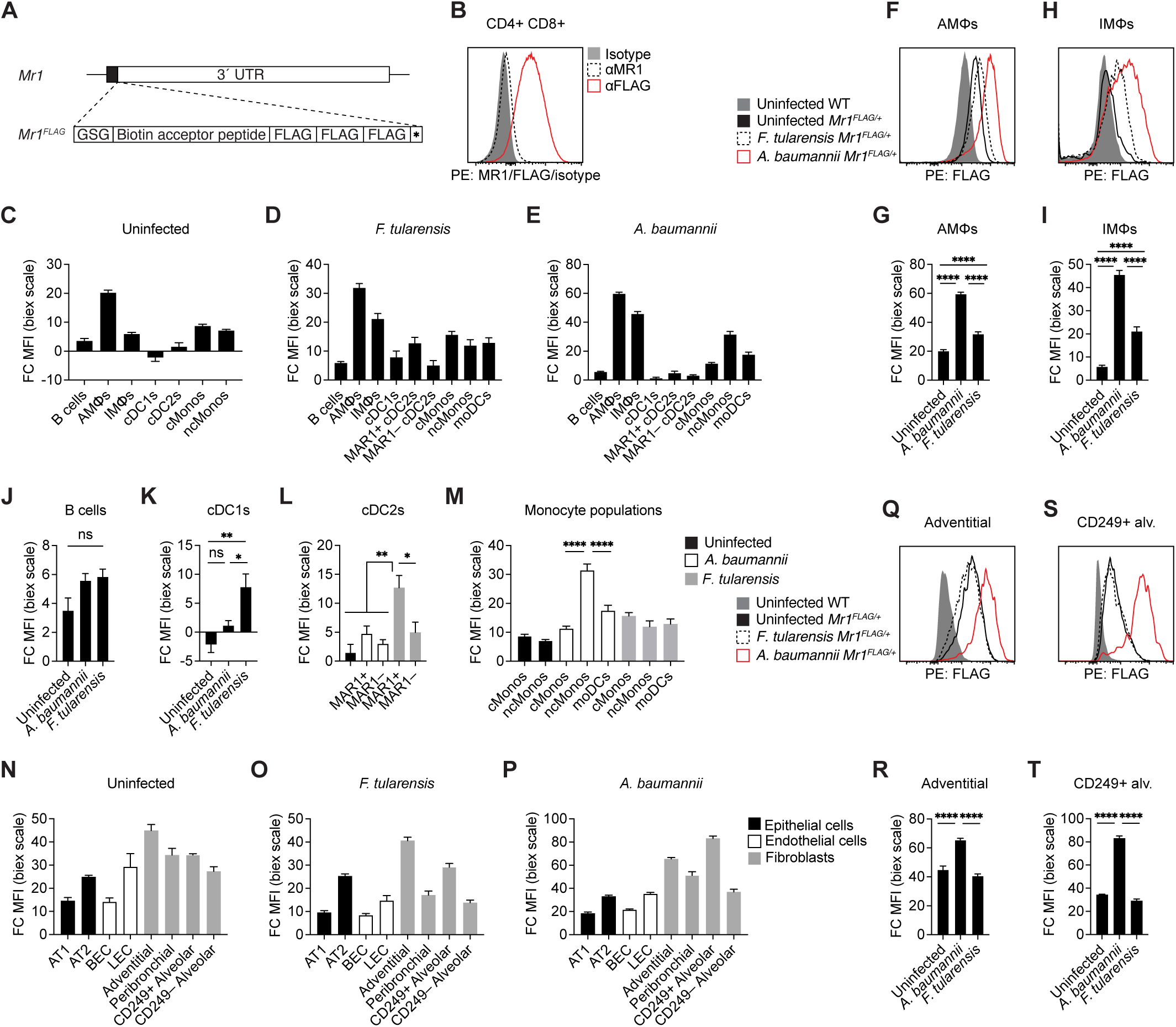
MR1 protein is broadly expressed and increased during infection. (A) Schematic for insertion of a GSG linker, biotin acceptor peptide, and 3 tandem FLAG tags following the translated portion of the 6^th^ exon of *Mr1*, prior to the STOP codon and 3′ UTR. (B) Representative flow staining of anti-FLAG and anti-MR1 on CD4^+^ CD8^+^ thymocytes. (C-E) Fold change in MFI (biexponential scale) compared to FLAG negative control for various hematopoietic subsets for uninfected mice (C), 4 DPI *A. baumannii* (D), or 4 DPI *F. tularensis* (E) (n=5-7/group). (F-I) Representative flow (F, H) and summary plots (G, I) of FLAG staining across conditions for AMΦs (F, G) and IMΦs (H, I). (J-M) Fold change in MFI (biexponential scale) compared to FLAG negative control for B cells (J), cDC1s (K), cDC2s separated by MAR1 staining (L) and monocyte populations (M) in uninfected mice, 4 DPI *A. baumannii*, or 4 DPI *F. tularensis*. (N-P) Fold change in MFI (biexponential scale) compared to FLAG negative control for various stromal subsets for uninfected mice (N), 4 DPI *A. baumannii* (O), or 4 DPI *F. tularensis* (P) (n=5-7/group). (Q-T) Representative flow (Q, S) and summary plots (R, T) of FLAG staining across conditions for adventitial fibroblasts (Q, R), and CD249^+^ alveolar fibroblasts (S, T) (n=5-7/group). Data representative of two or more independent experiments. Graphs indicate means ±SEM. One-way ANOVA (G, I, J-M, R, T) was performed. Statistics: ns, not significant (*P* > 0.05); \**P* < 0.05; \*\**P* < 0.01; \*\*\**P* < 0.001, \*\*\*\**P* < 0.0001.

### MR1 presentation is mediated by distinct APCs

MAIT cells respond to pathogens via TCR-dependent and TCR-independent activation.^8, 56^ Viruses stimulate human and murine MAIT cells independently of MR1 and cytokines are sufficient to induce activation *in vitro*.^57–60^ Therefore, we determined whether the MAIT cell response to bacterial pneumonia requires MR1 presentation. Since MAIT cells are selected on MR1 during their development,^10, 61^ we bred *Mr1^f/f^* conditional knockout mice to the *Rosa26-CreER^T2^* strain, where the ubiquitously expressed *Rosa26* drives expression of a *Cre* gene fused to the estrogen receptor gene *Esr1*, which retains the Cre recombinase in cytoplasm and prevents recombination until tamoxifen is administered. To ablate MR1 expression, we administered tamoxifen intraperitoneally (IP) to *Mr1^f/f^ Rosa26-CreER^T2^* and *Mr1^f/f^* mice prior to RPh inoculation with either *F. tularensis* or *A. baumannii*, the latter occurring 8 days later because tamoxifen metabolites inhibit *A. baumannii* growth **(Fig. 4A)**.^62^ While the number and frequency of MAIT cells in uninfected *Mr1^f/f^ Rosa26-CreER^T2^* mice and *Mr1^f/f^* littermates was comparable, ablation of MR1 severely inhibited the MAIT cell response to both pathogens **(Fig. 4B-C)**, indicating that MR1 presentation is necessary.

**Figure 4.**
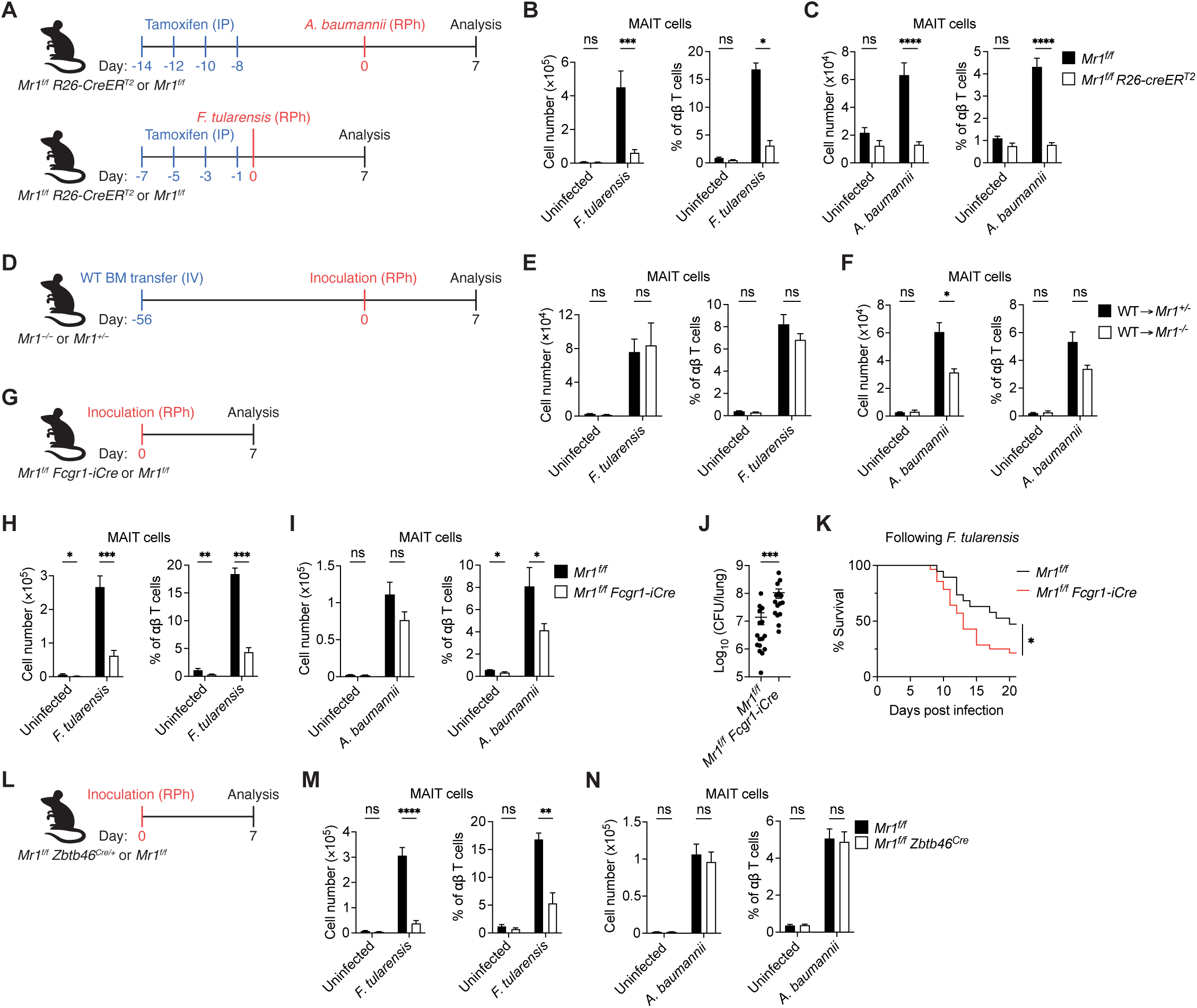
MR1 presentation is mediated by distinct APCs. (A-C) Schematic (A), number, and frequency of MAIT cells from the lungs of *Mr1^f/f^* and *Mr1^f/f^ Rosa26-creER^T2^* mice either uninfected or 7 DPI *F. tularensis* (B, n=7-17/group) or *A. baumannii* (C, n=4-19/group). (D-F) Schematic (D), number, and frequency of MAIT cells from the lungs of WT*→ Mr^+/−^* and WT *→ Mr1^−/−^*chimeric mice either uninfected or 7 DPI *F. tularensis* (E, n=6-7/group) or 7 DPI *A. baumannii* (F, n=3-4/group). (G-I) Schematic (G), number, and frequency of MAIT cells from the lungs of *Mr1^f/f^* and *Mr1^f/f^ Fcgr1-iCre* mice either uninfected or 7 DPI *F. tularensis* (H, n=4-9/group) or 7 DPI *A. baumannii* (I, n=4-11/group). (J) Bacterial load in the lungs of *Mr1^f/f^* and *Mr1^f/f^ Fcgr1-iCre* mice 10 DPI with 2×10^2^ CFU inoculum of *F. tularensis* (n=15-16/group). (K) Kaplan-Meier curve showing the survival of *Mr1^f/f^* and *Mr1^f/f^ Fcgr1-iCre* inoculated with 5×10^2^ CFU *F. tularensis* (n=19-27/group). (L-N) Schematic (L), number, and frequency of MAIT cells from the lungs of *Mr1^f/f^* and *Mr1^f/f^ Zbtb46^Cre^*mice either uninfected or 7 DPI *F. tularensis* (n=4-9/group) or 7 DPI *A. baumannii* (3-18/group). Data is representative of two or more independent experiments. Group numbers after each infection condition are inclusive of both uninfected and infected conditions as different uninfected mice were used in comparison to each infection model. Graphs indicate means ±SEM. Mann-Whitney U test (B, C, H, I, J, M, N), Mantel-Cox (K), and unpaired Student’s *t*-test (E, F) was performed. Statistics: ns, not significant (*P* > 0.05); \**P* < 0.05; \*\**P* < 0.01; \*\*\**P* < 0.001, \*\*\*\**P* < 0.0001.

To establish whether MR1 presentation is mediated by hematopoietic or non-hematopoietic cells, we generated bone marrow chimeras. Because MR1-mediated selection of MAIT cells occurs on DP thymocytes,^10, 61^ we transferred MR1-sufficient CD45.1^+^ WT bone marrow into irradiated CD45.2^+^ *Mr1^−/−^* and *Mr1^+/−^* recipients **(Fig. 4D)**. The resulting WT → *Mr1^−/−^* chimeras lacked MR1 presentation by non-hematopoietic cells, while WT → *Mr1^+/−^* chimeras retained MR1 presentation by both hematopoietic and non-hematopoietic cells. 8 weeks following the CD45.1^+^ bone marrow reconstitution, we verified that greater than 98% of pulmonary non-lymphocytes were CD45.1^+^ and subsequently inoculated with either *A. baumannii* or *F. tularensis* **(Fig. S4A)**. 7 days post-infection, the MAIT cell response to *A. baumannii* was partially reduced in WT → *Mr1^−/−^* chimeras compared to WT → *Mr1^+/−^* chimeras, while the MAIT cell response to *F. tularensis* was consistent in both **(Fig. 4E-F)**. These data indicate that hematopoietic cells mediate MR1 presentation during *F. tularensis* pneumonia, but both hematopoietic and non-hematopoietic cells present MR1 during *A. baumannii* infection.

Since MR1 was most highly expressed by macrophages among hematopoietic cells **(Fig. 3C-E)**, we interrogated whether these cells present microbial metabolites by crossing the *Mr1^f/f^* conditional knockout to a transgenic that expresses a codon-improved Cre (iCre) driven by the Fc receptor, IgG, high affinity I promoter (*Fcgr1-iCre*), which encodes the protein CD64.^63^ We verified specificity of iCre expression using the *Rosa26-LSL-eYFP* strain, which reports enhanced yellow fluorescent protein (eYFP) following excision of a *loxP*-flanked transcriptional STOP sequence (LSL). In *Mr1^f/f^ Fcgr1-iCre Rosa26-LSL-eYFP* mice, eYFP was expressed by ∼95% AMΦs and ∼45% of IMΦs, but we also observed substantial eYFP expression by monocytes, moDCs, and cDC2s **(Fig. S4B)**, which has been reported.^63^ Alternative Cre strains driven by the promoters of lysozyme 2 (*Lyz2^Cre^*) and colony-stimulating factor 1 receptor (*Csf1r-iCre*) are also expressed by monocytes and most cDCs, in addition to epithelial and lymphoid cells, respectively.^64^ Prior to infection **(Fig. 4G)**, *Mr1^f/f^ Fcgr1-iCre* mice exhibited fewer pulmonary MAIT cells than *Mr1^f/f^* animals **(Fig. 4H-I)**, indicating that MR1 presentation by cells that express or have expressed CD64 is necessary for the recruitment and/or maintenance of MAIT cells within the lungs. Following *F. tularensis* inoculation, *Mr1^f/f^ Fcgr1-iCre* mice accumulated substantially fewer pulmonary MAIT cells than *Mr1^f/f^* animals, but their MAIT cell frequency was only slightly reduced during *A. baumannii* infection **(Fig. 4H-I)**. The diminished MAIT cell response to *F. tularensis* impaired bacterial clearance by *Mr1^f/f^ Fcgr1-iCre* mice and decreased their survival **(Fig. 4J-K)**. These results demonstrate that MR1 presentation by hematopoietic cells induces a protective MAIT cell response to *F. tularensis*.

Since ∼50% of cDC2s were eYFP^+^ in *Mr1^f/f^ Fcgr1-iCre Rosa26-LSL-eYFP* mice **(Fig. S4B)**, we determined whether cDCs mediate MR1 presentation using a Cre strain driven by the zinc finger and BTB domain-containing 46 promoter (*Zbtb46^Cre^*).^65^ Due to expression of *Zbtb46* at the pre-DC stage, Cre expression is limited to cDCs, with minimal excision in plasmacytoid DCs and macrophages.^65^ Using *Mr1^f/f^ Zbtb46^Cre^ Rosa26-LSL-eYFP* mice, we confirmed that eYFP was expressed by cDC1 and cDC2s, with little expression by macrophages, monocytes, and moDCs **(Fig. S4C)**. In uninfected *Mr1^f/f^ Zbtb46^Cre^* and *Mr1^f/f^*mice **(Fig. 4L)**, the abundance of pulmonary MAIT cells was similar **(Fig. 4M-N)**, indicating that MAIT cells do not require MR1 presentation by cDCs at homeostasis. However, the MAIT cell response to *F. tularensis* was abrogated in *Mr1^f/f^ Zbtb46^Cre^* mice, while the response to *A. baumannii* was unaltered **(Fig. 4M-N)**. Together with the previous results, these data suggest that non-hematopoietic cells and redundant hematopoietic populations present *A. baumannii* metabolites, while MR1 presentation of *F. tularensis* metabolites is primarily mediated by cDCs.

### MAIT cell responses originate in different locations

MAIT cell responses to commensal and pathogenic bacteria have been reported to occur in barrier tissues and their draining lymph nodes.^10, 20, 66^ Given the disparate roles for non-hematopoietic cells during *A. baumannii* and *F. tularensis* infections, we interrogated whether MAIT cell activation occurs in the lungs or mediastinal lymph nodes (meLNs) by administering the sphingosine 1-phosphate receptor (S1PR) agonist FTY720, which induces S1PR internalization and prevents lymphocyte egress from secondary lymphoid organs **(Fig. 5A)**.^67, 68^ FTY720 depleted CD4^+^ and CD8^+^ T cells from circulation and abrogated their accumulation within the lungs during both infections **(Fig. S5A-C)**. During *F. tularensis* infection, FTY720 decreased pulmonary MAIT cells by ∼80% **(Fig. 5B)**, suggesting their response originates within the meLNs. Conversely, FTY720 only diminished MAIT cells by ∼50% following *A. baumannii* inoculation **(Fig. 5C)**, indicating that the MAIT cell response to this pathogen can occur within the lungs or meLNs. To determine where MAIT cell proliferation commences following *F. tularensis*, we measured expression of the nuclear protein Ki67, which is expressed during cell division. Although there were few Ki67^+^ MAIT cells prior to infection, *F. tularensis* induced MAIT cell proliferation 4 days post-inoculation, with a higher frequency of Ki67^+^ MAIT cells in the meLNs compared to the lungs **(Fig. 5D)**, suggesting that MAIT cells divide in the meLNs and subsequently traffic to the lungs. While ∼70% of pulmonary MAIT cells were labeled by an IV-injected anti-CD45.2 antibody in naïve animals, inoculation with *F. tularensis* decreased the frequency of vascular MAIT cells to less than 10% within 4 days **(Fig. 5E)**, signifying rapid entry into the lung parenchyma. Together, these results demonstrate that MAIT cells respond to *A. baumannii* in the lungs and meLNs, while they proliferate in meLNs during *F. tularensis* infection and subsequently transit to the lung parenchyma.

**Figure 5.**
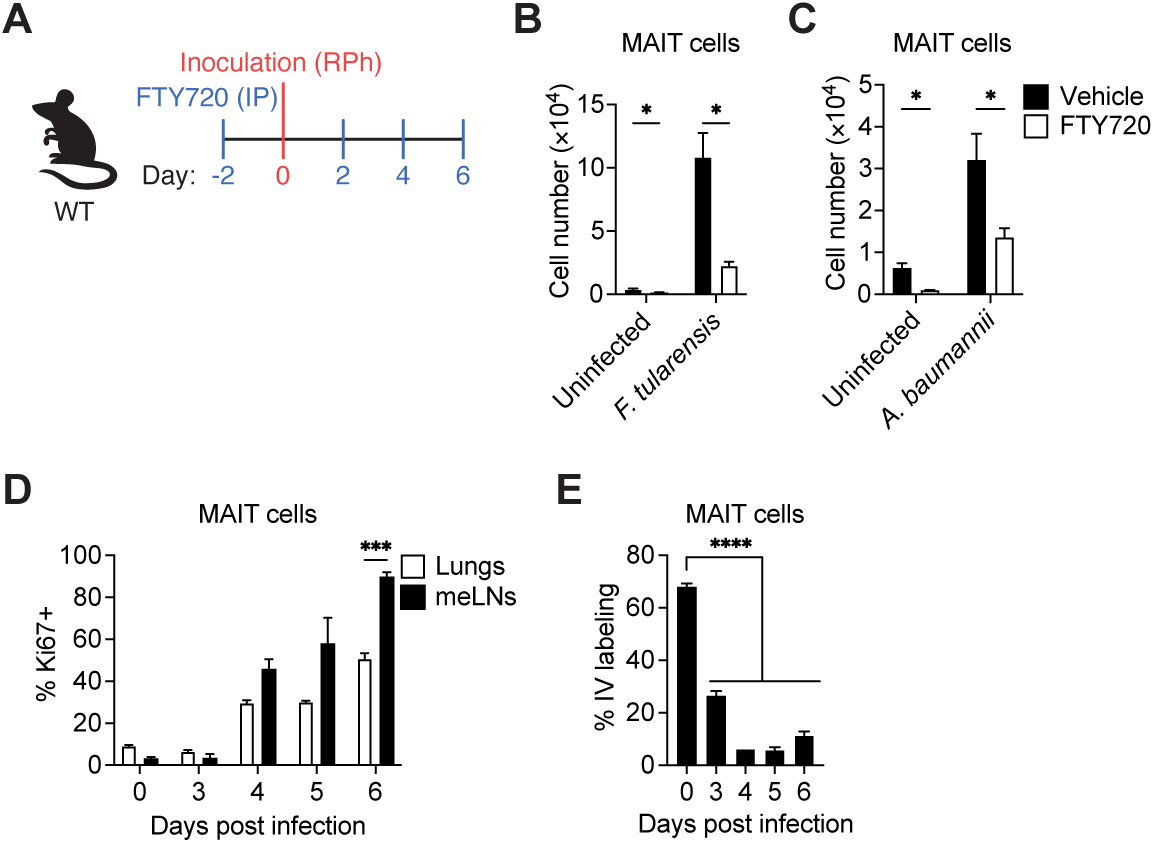
MAIT cell responses originate in different locations. (A) Schematic of FTY720 treatment regimen and infection inoculum schedule. (B) Number of MAIT cells in the lungs of mice treated with vehicle or FTY720 either uninfected or 7 DPI *F. tularensis* (n=5/group). (C) Number of MAIT cells in the lungs of mice treated with vehicle or FTY720 either uninfected or 7 DPI *A. baumannii* (n=5-6/group). (D) Ki-67 levels from MAIT cells in the lungs and meLNs on 3-6 DPI *F. tularensis* (n=2-4/group). (E) Frequency of αCD45.2^+^ labelling in the lungs of mice 3-6 DPI. Infected mice were IV injected with αCD45.2^+^ three minutes prior to tissue collection (n=1-4/group). Data is representative of two independent experiments (B, C). Graphs indicate means ±SEM. Unpaired Student’s *t*-test (B-D) and one-way ANOVA (E) was performed. Statistics: ns, not significant (*P* > 0.05); \**P* < 0.05; \*\**P* < 0.01; \*\*\**P* < 0.001, \*\*\*\**P* < 0.0001.

### Inflammatory cDC2s present *F. tularensis* metabolites

AMΦs account for 70% of the cells infected with *F. tularensis* LVS 1 day following pulmonary inoculation.^69^ To determine whether AMΦs mediate the MAIT cell response to *F. tularensis* or *A. baumannii*, we administered liposomes containing clodronate or phosphate-buffered saline (PBS) intranasally (IN) 2 days prior to inoculation and daily thereafter **(Fig. 6A)**. We confirmed that clodronate liposomes depleted AMΦs without altering the abundance of other myeloid cells during the infection **(Fig. S6A-F)**. Clodronate treatment reduced the number of MAIT cells that responded to *F. tularensis*, but not *A. baumannii* **(Fig. 6B-C)**. However, AMΦs provide an important replicative niche for *F. tularensis* and their depletion with clodronate liposomes has been shown to decrease *F. tularensis* CFUs during pulmonary infection.^70^ We confirmed that clodronate-mediated depletion of AMΦs diminished growth of the *F. tularensis* inoculum **(Fig. 6D-E)**. Given that the MAIT cell response to *F. tularensis* was abrogated in *Mr1^f/f^ Zbtb46^Cre^* mice and the *Zbtb46^Cre^* allele was not expressed by AMΦs **(Figs. 4M & S4C)**, we conclude that AMΦs facilitate bacterial replication rather than MR1 presentation during *F. tularensis* infection and are not required for the MAIT cell response to *A. baumannii*, perhaps due to redundancy with other APCs.

**Figure 6.**
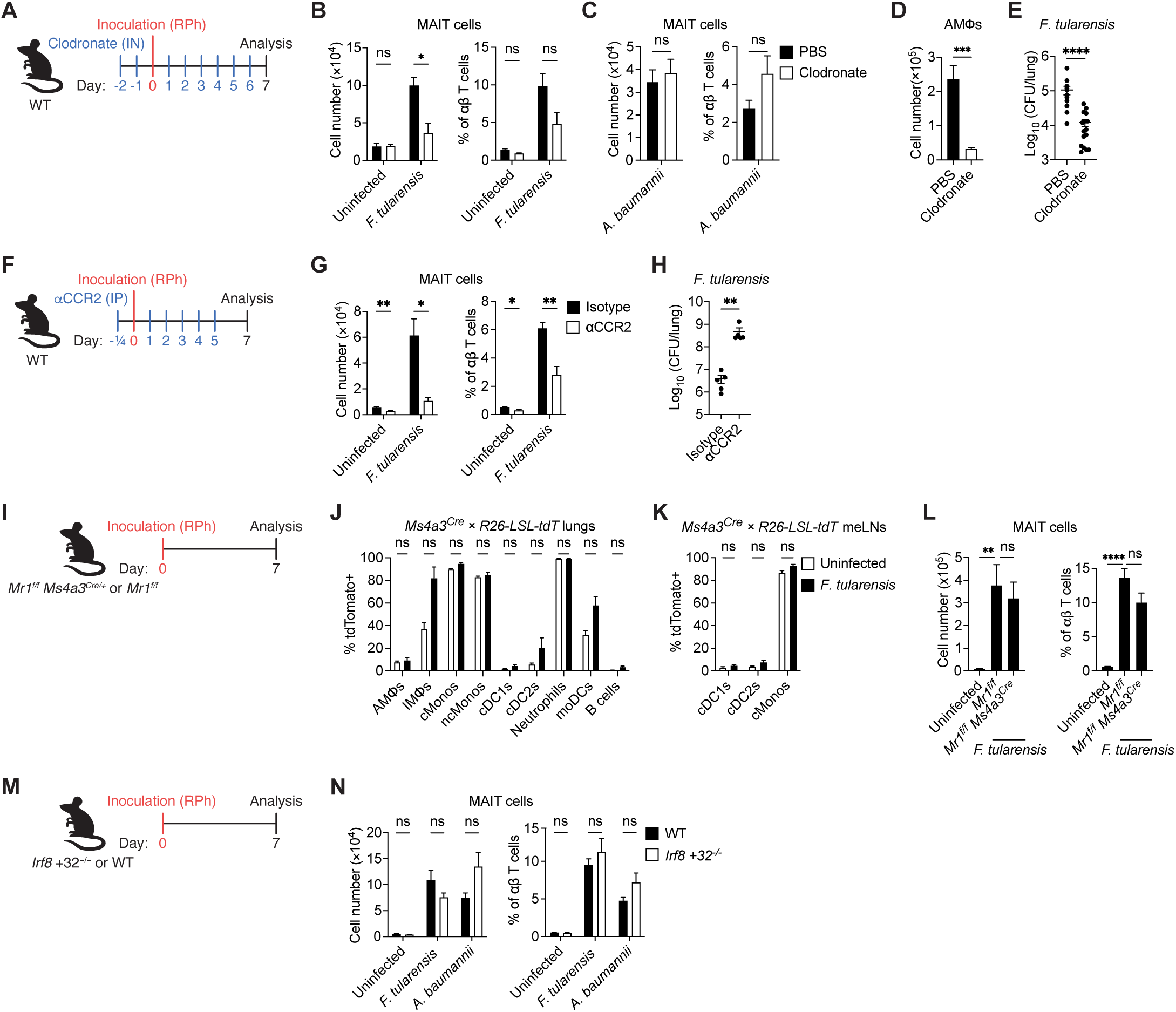
Inflammatory cDC2s present *F. tularensis* metabolites. (A) Schematic of clodronate liposome administration and infection. (B) Number and frequency of MAIT cells in the lungs of clodronate or PBS liposome treated mice 7 DPI *F. tularensis* (n=4-6/group). (C) Number and frequency of MAIT cells in the lungs of clodronate or PBS liposome treated mice 7 DPI *A. baumannii* (n=4-6/group). (D) Number of AMΦ after 2 days of daily treatment with PBS or clodronate liposomes prior to infection (n=9/group). (E) Bacterial load in the lungs 1 DPI of 2×10^3^ CFU *F. tularensis* after clodronate or PBS liposome treatment on -2 and -1 DPI (n=14-16/group). (F) Schematic of αCCR2 administration and infection. (G) Number and frequency of MAIT cells from the lungs of mice after IP treatment of αCCR2 or isotype control either uninfected or 7 DPI *F. tularensis* (n=5/group). (H) Bacterial load in the lungs of αCCR2 and isotype treated mice 7 DPI with 2×10^3^ CFU *F. tularensis* (n=5/group). (I) Schematic of *F. tularensis* infection and analysis in *Mr1^f/f^ Ms4a3cre* mice. (J) tdTomato expression in the lungs of uninfected *Ms4a3^Cre^* × *Rosa26-LSL-tdTomato* mice or inoculated with 2×10^3^ CFU *F. tularensis (*n=6/group). (K) tdTomato expression in the meLNs of uninfected *Ms4a3^Cre^* × *Rosa26-LSL-tdTomato* mice or inoculated with 2×10^3^ CFU *F. tularensis (*n=6/group). (L) Number and frequency of MAIT cells from the lungs of *Mr1^f/f^* and *Mr1^f/f^ Ms4a3^Cre^* mice 7 DPI *F. tularensis* (n=5-7/group). (M) Schematic of infection and analysis in *Irf8 +32^-/-^* mice (N) Number and frequency of MAIT cells from the lungs of WT and *Irf8 +32^-/-^* mice either uninfected or 7 DPI *F. tularensis* (n=6-11/group) or 7 DPI *A. baumannii* (n=6-11/group). Data is representative of two or more independent experiments. Graphs indicate means ±SEM. Unpaired Student’s *t*-test (D, C, G, K) unpaired Mann-Whitney U test (B, E, H, J, N), or one-way ANOVA (L) was performed. Statistics: ns, not significant (*P* > 0.05); \**P* < 0.05; \*\**P* < 0.01; \*\*\**P* < 0.001, \*\*\*\**P* < 0.0001.

In addition to replicating in macrophages, pulmonary *F. tularensis* LVS also infects monocytes and DCs.^69^ To interrogate if monocytes or monocyte-derived cells are necessary for the MAIT cell response to *F. tularensis*, we utilized a depletion antibody specific for C-C motif chemokine receptor 2 (CCR2), which is highly expressed by classical monocytes.^71^ However, CCR2 is also expressed by some T cells,^71^ so we measured the expression on the MAIT1 cells that preferentially respond to *F. tularensis* **(Fig. S6G)**. While pulmonary MAIT1 cells did not express CCR2 before or during *F. tularensis* infection, MAIT17 cells were CCR2^+^ in uninfected animals, although the few *F. tularensis*-responsive MAIT17 cells lacked CCR2 expression **(Fig. S6H-I)**. Thus, anti-CCR2 mediated depletion of monocytes is a viable approach for establishing their role during *F. tularensis* infection. To deplete monocytes, we injected anti-CCR2 (clone MC-21) IP 5 hours prior to inoculation with *F. tularensis* and daily thereafter until day 5 **(Fig. 6F)**, at which point neutralizing antibodies render anti-CCR2 ineffective. Classical monocytes were abrogated by anti-CCR2, which also depleted cDC1s, cDC2s, and IMΦs **(Fig. S6J-O)**, the latter of which is repopulated by classical monocytes and expresses intermediate levels of CCR2.^72, 73^ While depletion of cDCs was unexpected, their pre-cDC precursors express CCR2, which facilitates their recruitment to the lungs during infection.^74^ Anti-CCR2 depletion abolished the MAIT cell response to *F. tularensis* and impaired bacterial clearance **(Fig. 6G-H)**, indicating that monocytes, monocyte-derived cells, and/or cDCs are necessary for MAIT cell-mediated immunity.

Since monocyte-derived DCs (moDCs) have been shown to uptake *F. tularensis* bacteria,^44^ we determined whether they present *F. tularensis* metabolites using a Cre strain driven by the membrane-spanning 4-domains, subfamily A, member 3 promoter (*Ms4a3^Cre^*). *Ms4a3* is transiently expressed by the granulocyte-monocyte progenitor and the subsequent granulocyte progenitor and common monocyte progenitor, which limits Cre excision to monocytes, monocyte-derived cells, and granulocytes.^75^ Using *Mr1^f/f^ Ms4a3^Cre^ Rosa26-LSL-tdTomato* mice, we verified that tdTomato was expressed by pulmonary monocytes, monocyte-derived cells, and granulocytes, but not cDCs nor AMΦs **(Fig. S6P)**. In the meLNs, cDCs were tdTomato^−^ **(Fig. S6Q)**, confirming they were not monocyte-derived. Following *F. tularensis* inoculation **(Fig. 6I)**, the specificity of Cre expression was largely unchanged, except for the increased frequency of tdTomato^+^ moDCs and IMΦs that arise from monocytes **(Fig. 6J-K)**. The MAIT cell response to *F. tularensis* was comparable between *Mr1^f/f^ Ms4a3^Cre^* and *Mr1^f/f^* mice **(Fig. 6L)**, indicating that MR1 presentation by monocytes, monocyte-derived cells, and granulocytes is not necessary during this infection.

Since the MAIT cell response to *F. tularensis* was diminished by anti-CCR2 depletion of monocytes and pre-cDCs but retained in *Mr1^f/f^ Ms4a3^Cre^* mice that lack MR1 expression on monocyte-derived cells, we conclude that MR1 presentation depends on cells derived from CCR2^+^ pre-cDCs. The abrogated MAIT cell response in *Mr1^f/f^ Fcgr1-iCre* and *Mr1^f/f^ Zbtb46^Cre^* mice suggests that *F. tularensis* metabolites are presented by inf-cDC2s, which express *Fcgr1*.^54^ Therefore, we determined whether MAIT cell responses are dependent on cDC1s, which require interferon regulatory factor 8 (*Irf8*) for their development and are absent from mice that lack the +32 kilobase *Irf8* enhancer (*Irf8 +32^−/−^*).^76^ We confirmed that cDC1s were absent from *Irf8 +32^−/−^*mice, which retained other myeloid populations **(Fig. S6R-W)**. The number and frequency of MAIT cells was comparable between *Irf8 +32^−/−^* and WT mice prior to infection and following inoculation with *F. tularensis* or *A. baumannii* **(Fig. 6M-N)**, indicating that cDC1s do not determine MAIT cell abundance at homeostasis nor during infection. These results suggest that inf-cDC2s mediate MR1 presentation of metabolites from the intracellular bacterial pathogen *F. tularensis*.

## Discussion

Our work identified *A. baumannii* as the first extracellular pathogen that induces robust MAIT cell proliferation. Conservation of riboflavin synthesis among *A. baumannii* isolates permitted MAIT cell responses to diverse strains of this pathogen in an MR1-dependent manner. MAIT17 cells were the primary IL-17A producers during *A. baumannii* pneumonia and transcriptionally distinct from the MAIT1 cells that responded to the intracellular pathogen *F. tularensis*. The MAIT17 cells expressed transcriptional signatures associated with tissue residency and repair while the MAIT1 cells transcribed proliferation genes. Utilizing a novel *Mr1^FLAG^* reporter strain, we determined that MR1 protein is most highly expressed by AMΦs and non-hematopoietic cells at homeostasis and *A. baumannii* infection enhanced expression on these populations as well as IMΦs. While microbial metabolites are known to facilitate trafficking of MR1 to the plasma membrane,^26, 27^ increased intracellular MR1 expression suggests transcriptional and/or translational regulation in response to *A. baumannii*, similar to interferon-induced MHC-I transcription.^77^ The partial reduction of MAIT cells observed in *Mr1^f/f^ Fcgr1-iCre* mice and WT → *Mr1^−/−^* bone marrow chimeras following *A. baumannii* infection indicates that both CD64^+^ hematopoietic and non-hematopoietic cells present metabolites via MR1. However, clodronate-mediated depletion of AMΦs and deletion of MR1 from cDCs in *Mr1^f/f^ Zbtb46^Cre^* mice were insufficient to diminish the MAIT cell response to *A. baumannii*, so we conclude that multiple redundant hematopoietic populations present metabolites from this extracellular pathogen. Previous studies have shown that epithelial cells can activate MAIT cells by presenting metabolites derived from *M. tuberculosis*, *Shigella flexneri*, and *S.* Typhimurium.^78–80^ While alveolar epithelial cells express MR1 protein, we observed significantly greater expression by fibroblasts. As observed for AMΦs and IMΦs, *A. baumannii* infection enhanced their expression of MR1, with the largest increases by adventitial and CD249^+^ alveolar fibroblasts. These data suggest that fibroblasts may present metabolites from extracellular bacteria, warranting further investigation.

While non-hematopoietic cells present *A. baumannii* metabolites, WT → *Mr1^−/−^* bone marrow chimeras indicated that their expression of MR1 is not required for the MAIT cell response to *F. tularensis*, which concurs with the necessity for hematopoietic MR1 expression during respiratory infection with another intracellular pathogen, *L. longbeachae*.^80^ Although *F. tularensis* subverts CD4^+^ T cell activation by inducing ubiquitination and degradation of MHC-II,^81, 82^ MR1 levels were sustained during infection, with substantially greater expression by cDC2s and IMΦs. *F. tularensis* replication within AMΦs causes apoptosis of infected cells,^83^ which may have diminished their contribution to metabolite presentation despite their high expression of MR1. *L. longbeachae* also replicates in AMΦs and results in their depletion,^84^ so other intracellular respiratory pathogens may hinder MR1 presentation by AMΦs as well. The decreased MAIT cell accumulation in *Mr1^f/f^ Zbtb46^Cre^* mice following *F. tularensis* suggested a requirement for MR1 presentation by cDCs, while the sustained response in *Irf8 +32^−/−^* and *Mr1^f/f^ Ms4a3^Cre^* animals precluded a role for cDC1s and monocyte-derived cells, respectively. Furthermore, the MAIT cell response to *F. tularensis* was impaired by anti-CCR2 depletion of monocytes and pre-cDC precursors as well as in *Mr1^f/f^ Fcgr1-iCre* mice. From these results we conclude that MR1 presentation of *F. tularensis* metabolites is mediated by cDCs that express CD64, implying inf-cDC2s.^54^ These APCs stimulate viral-specific CD4^+^ and CD8^+^ T cells to a greater degree than cDC2s and moDCs,^54^ demonstrating their capacity for antigen presentation. While MR1 presentation by inf-cDC2s has not been demonstrated previously, DCs were ∼1,000-fold more efficient than monocyte, B cell, and bronchial epithelial cell lines at activating an MR1-restricted T cell clone *in vitro*.^49^ Although the MAIT cell response to *F. tularensis* was sustained in *Mr1^f/f^ Ms4a3^Cre^ Rosa26-LSL-tdTomato* mice where a substantial portion of IMΦs exhibited *Ms4a3^Cre^* activity, these cells may also present *F. tularensis* metabolites since they expanded during infection and highly expressed MR1. Murine models that only ablate IMΦs or their presentation of MR1 are needed to determine whether they contribute to the MAIT cell response.

DCs have been implicated in the presentation of microbial antigens to other unconventional T cells. Deletion of CD1d from Langerin^+^ cDC1s decreased IL-17A production by iNKT cells during *S. pneumoniae* infection and inhibited bacterial clearance from the lungs.^85^ Loss of CD1d expression on CD11c^+^ DCs and macrophages prevented iNKT responses to a microbially-derived glycolipid and altered composition of the intestinal microbiota at homeostasis.^86, 87^ Given that *Mr1^−/−^* and WT mice retain distinct intestinal microbes following cohousing,^88^ elucidating which APCs present commensal-derived metabolites is of interest. The reduction of pulmonary MAIT cells in naïve *Mr1^f/f^* mice bearing the broadly expressed *Fcgr1-iCre*, but not the predominantly cDC-specific *Zbtb46^Cre^*or monocyte/granulocyte-specific *Ms4a3^Cre^* alleles, suggests that MAIT cell development and/or maintenance within the lungs is mediated by MR1-expressing macrophages or redundant hematopoietic populations. TCR-mediated activation of MAIT cells induces their expression of tissue repair genes and ameliorates experimental autoimmune encephalomyelitis,^57, 89^ although the cells that present MR1 in the context of non-microbial responses remain to be determined.

Despite broad expression of MR1, our findings demonstrate that microbial tropism dictates which cells mediate presentation of metabolites. Consequently, targeted delivery of MR1 ligands to the appropriate APCs may enhance the responses of MAIT cells and other MR1-resticted T cells.

## Acknowledgements

The authors acknowledge the Scripps Research Department of Animal Resources, Flow Cytometry Core, and Genomics Core and thank Dr. Olivier Lantz for the WT3-MR1 cells and 8D12 MAIT hybridoma,^40, 41^ Dr. Luc Teyton for the DC2.4 and DC3.2 cell lines, Dr. Bernard Malissen for the *Fcgr1-iCre-mTFP1* strain,^63^ and the NIH Tetramer Core Facility for mCD1d and mMR1 tetramers. This research was supported by the National Institutes of Health (K22AI146217, R21AI171697, & R35GM151347 to M.G.C.), the Natural Sciences and Engineering Research Council of Canada (Doctoral Postgraduate Scholarship to D.H.), the National Science Foundation (Graduate Research Fellowship to G.L.), and Scripps Research.

## Author contributions

Conceptualization: D.H., J.M., and M.G.C.; Methodology: D.H., J.M., and M.G.C.; Investigation: D.H., J.M., and G.L.; Analysis: D.H., J.M., N.A., and M.G.C.; Writing – Original Draft: D.H., J.M., and M.G.C.; Writing – Review & Editing: D.H., J.M., and M.G.C.; Visualization: D.H., J.M., and M.G.C.; Supervision: M.G.C.; Funding Acquisition: M.G.C., D.H., J.M., and G.L.; Resources: A.J.P., J.A., and M.M.

## Methods

### Experimental model and subject details

WT (C57BL/6J), CD45.1 (C57BL/6J-Ptprc^em6Lutzy^/J), *Irf8+32^-/-^* (C57BL/6-Rr172^em1Kmm^/J), *Ms4a3^Cre^*(C57BL/6J-Ms4a3^em2(cre)Fgnx^/J), *Fcgr1-iCre* (B6(129S4)-Fcgr1^em1.1(icre)Moli^/J), *Zbtb46^Cre^* (B6.Cg-Zbtb46^tm3.1(cre)Mnz^/J), *Rosa26-CreER^T2^*(B6.129-Gt(ROSA)26Sor^tm1(cre/ERT2)Tyj^/J), *Rosa26-LSL-eYFP* (B6.129X1-Gt(ROSA)26Sor^tm1(EYFP)Cos^/J), *Rosa26-LSL-tdTomato* (B6.Cg-Gt(ROSA)26Sor^tm^^14(CAG-tdTomato)Hze^/J) were obtained from the Jackson Laboratory. *Mr1^-/-^* (*Mr1^tm1Gfn^)* mice were generated by Dr. Susan Gilfillan.^90^ *Mr1^f/f^*mice were generated as previously described.^10^ *Mr1^f/f^*mice were crossed to *Zbtb46^Cre^*, *Ms4a3^Cre^, or Fcgr1-iCre* (gift of Dr. Bernard Malissen) to generate *Mr1^f/f^* homozygous Cre heterozygous mice.^63, 65^ *Mr1^f/f^* × *Rosa26-creER^T2^* mice were generated and the inducible deletion of MR1 was performed with IP injection of tamoxifen. All experiments were age-matched between 7-9 weeks of age. To account for cage effect on immune cell populations (MAIT cell frequency from microbiota), litters were spread across different experimental groups.^10^ Mice were bred and cared for in a facility accredited by the American Association for the Accreditation of Laboratory Animal Care at the Scripps Research Institute. All experiments were conducted at Scripps Research in accordance with an Animal Safety Protocol approved by the Scripps Research Institutional Animal Care and Use Committee.

### Chimera generation

Recipient mice were subjected to two rounds of 5.75Gy X-ray irradiation three hours apart. Donor bone marrow cells were isolated via centrifugation as previously described.^91^ Donor bone marrow cells were resuspended to 10^8^ cells/mL in PBS and 100uL was injected to host mice via retroorbital injection. After injection, mice were placed in sterile housing with TMS water. One week later, mice were removed from TMS water. Two days after antibiotic removal, microbiota was reconstituted as previously described.^92^ Cecal contents were harvested anaerobically and 100uL of 50mg/mL cecal contents in reduced PBS was given to host mice via oral gavage. Mice were rested for 6 weeks prior to blood check for bone marrow reconstitution and 8 weeks prior to infection.

### Bacterial culture

Cultures of *Francisella tularensis* subsp. *holarctica,* CDC Live Vaccine Strain (BEI Resources, NR-646) were grown in tryptic soy broth supplemented with 0.1g/100ml of cysteine (TSBC) at 37°C with shaking 220 rpm. OD/CFU curves were generated with an optimal linear regression identified between an OD of 0.5-0.8. To get an infectious inoculum, a single bacterial colony was grown in TSBC for 8-12 hours before back diluting to an OD of 0.01. This culture was grown for another 14-18 hours to reach an appropriate OD between 0.5-0.8. Culture was diluted in PBS to appropriate infectious dose. Prior to infection, inoculums were plated on cysteine heart agar plates enriched with 1% hemoglobin for 3 days to verify dosage.

Cultures of *A. baumannii* (ATCC 17978) were grown in lysogeny broth (LB) at 37°C with shaking 220 rpm. OD/CFU curves were generated with an optimal linear regression identified between an OD of 0.5-0.8. Overnight culture from a single colony was back diluted to 0.01 prior to infection. At proper OD, bacteria were concentrated by centrifugation at 4000rcf for 12 minutes. Bacteria was resuspended in PBS for instillation into mice. Prior to infection, inoculums were plated on LB agar for overnight growth to verify dosage.

### Inoculation of mice and survival analysis

Age- and sex-matched mice were anaesthetized with isoflurane and inoculated with 40ul of specified inoculum of *F. tularensis* LVS or *A. baumannii* RPh via reflexive aspiration. For model optimization and survival studies, weight was assessed daily and mice were considered moribund after a weight loss over 25%. *F. tularensis* inoculums were 2×10^3^ CFU for all experiments except for survival studies for which a dose 5×10^3^ CFU was used. *A. baumannii* inoculums were between 0.8-1×10^8^ CFU for all experiments.

### Tissue processing

Lungs were harvested in PBS on ice and subsequently finely minced in 2ml of digest media using razor blade. Digest media included supplemented RPMI (RPMI 1640 with 1 mM sodium pyruvate, 20mM glutaGRO, 1 mM non-essential amino acids, 50 mM β-mercaptoethanol, 20mM HEPES, 100 U/mL penicillin, 100 mg/mL streptomycin), 0.5 mg/mL DNase I, and 0.25 mg/ml Liberase TL. Minced lung in digest media was incubated for 45 min in a 37°C water bath and gently vortexed every 15 minutes. Digested lungs were dissociated through a 70μm cell strainer and washed with 5 ml of supplemented RPMI media containing 0.5mg/ml DNase and 3% fetal bovine serum (FBS). Single cell suspensions were centrifuged at 526rcf for 5 minutes and resuspended in 2 mL of 37% Percoll. After centrifugation and aspiration of Percoll supernatant, pellet was resuspended in 2mL RBC Lysis Buffer for 3 min at room temperature and stopped with 3mL of PBS. After centrifugation, pellets were resuspended in supplemented RPMI with 10% FBS (RPMI10) before processing for flow cytometry.

MeLNs were harvested PBS on ice and subsequently digested in 0.5 ml of supplemented RPMI with 0.5 mg/mL DNase I and 0.05 mg/ml Liberase TL for 20 minutes at 37°C in a 5% CO2 incubator. Digested meLNs were dissociated through a 70μm cell strainer and washed with 2 ml of 3% DNase. After centrifugation, pellets were resuspended in complete RPMI with 10% FBS before processing for flow cytometry.

For isolation of lung stromal cells, Dispase was added to digest media at 0.25U/mL.^93^ Percoll supernatant was kept and used for downstream stromal staining.

### Flow cytometry

Single cell suspensions were stained in viability stain in PBS for 20 minutes on ice in the dark. Extracellular stain was completed in RPMI10 for 1 hour at room temperature (lymphoid panel; for tetramer stain) or 30 minutes on ice (myeloid, non-hematopoietic panels) in the dark. MAR-1 secondary antibody myeloid staining was performed in PBS for 20 minutes on ice in the dark following extracellular stain due to presence of biotin in RPMI10. Cells were fixed in fixative from eBioscience FoxP3/Transcription Factor kit for 1 hour on ice in the dark. Intracellular staining was completed overnight in permeabilization buffer from eBioscience FoxP3/Transcription Factor kit. Cells were resuspended in FACS buffer (PBS+2% FBS) prior to collection on Cytek Aurora.

In the case of fluorescent protein detection, BD Cytofix for 30 minutes on ice was used instead of eBioscience FoxP3/Transcription Factor kit to prevent quenching of signal. Following BD Cytofix, cells were washed in FACS buffer prior to collection on Cytek Aurora.

Lymphoid and myeloid cells were gated as shown **(Figs. S1A, S1C-D, & S3C)** and non-hematopoietic cells were identified as previously described.^55^

### *Ex vivo* restimulation

For *ex vivo* restimulation, single cell suspensions in RPMI10 with 1x PMA/ionomycin and 1x Brefeldin A were stimulated at 37°C in a 5% CO_2_ incubator for 2.5 hours. Flow cytometry staining was performed as described above with appropriate cytokines to detect restimulation of cells.

### Genome Computational Analysis

Nucleotide sequences of the following riboflavin synthesis genes were downloaded from the National Center for Biotechnology Information: *ribA* (NZ_CP045110.1:3590616-3591218), *ribB* (NZ_CP045110.1:c868072-867413), *ribD* (NZ_CP045110.1:301694-302779), *ribE* (NZ_CP045110.1:3901151-3901621), *ribH* (NC_016603.1:3072351-3072821), *pyrp2* (WP_001829328.1). These sequences were queried using BLASTn and tBLASTn for *pyrp2* search against the National Center for Biotechnology Information core_nt database using an e-value <1E^-10^ (version BLAST 2.15.0). Alignments that met this threshold were considered homologs. The percentage of strains containing riboflavin synthesis genes was determined by comparing alignments against the list of complete genomes for each of the following *Acinetobacter* species in the core_nt database: *A. baumannii* (txid: 470; n=965), *A. calcoaceticus* (txid: 471; n=6), *A. nosocomialis* (txid: 106654; n=22), *A. pittii* (txid: 48296; n=79).

### Evaluation of bacterial burden

Harvested lungs were chopped and dissociated through 70μm cell strainers and washed with 7 ml of supplemented RPMI media containing 0.5mg/ml DNase and 3% fetal bovine serum. Single cell suspensions were diluted in PBS prior to plating. CFU per lung were determined by colony enumeration 3 days after plating suspensions from *F. tularensis* LVS-infected lungs on CHA and 1 day after plating suspensions from *A. baumannii*-infected lungs on LB.

### Single cell RNA sequencing

MAIT cells were single cell sorted as TCRβ^+^, MR1:5-OP-RU tetramer^+^ from uninfected mice, mice 7 DPI *F. tularensis*, or mice 7 DPI *A. baumannii* on Thermo Fisher Scientific Bigfoot. Single cell RNAseq libraries were acquired utilizing 10x Genomics manufacturer recommendations by Scripps Research Genomics core. Prior to analysis, all three libraries were combined utilizing 10x Genomics Cell Ranger for a single analysis. The combined single cell RNA-seq dataset was processed, explored and visualized using Trailmaker^TM^ (https://app.trailmaker.parsebiosciences.com/; Parse Biosciences). Unfiltered count matrices were uploaded to Trailmaker, and background was removed by setting a minimum transcripts per cell threshold (threshold range: 1000 minimum). Dead or dying cells were removed by filtering barcodes with high mitochondrial content (10% cut-off). Outliers in the distribution of number of genes vs number of transcripts were removed by fitting a linear regression model (p-values between 0.000082 and 0.000660). Cells with a high probability of being doublets were filtered out using the scDblFinder method (threshold range: 50%). Cells with expression of *Adgre1, Itgam, Itgax, Fcgr1, Ms4a1, Mrc1, C1qc, Cd19, Cd79a* and those identified as non-classical monoyctes by Trailmaker^TM^ platform were filtered out (∼2% of total dataset). Data normalization, principal-component analysis (PCA) and data integration using Harmony were performed on data from high-quality cells. Clusters were identified using the Louvain method, and a Uniform Manifold Approximation and Projection (UMAP) embedding was calculated to visualize the results. Cluster-specific marker genes were identified by comparing cells of each cluster to all other cells using the “pRESTO” package implementation of the Wilcoxon rank-sum test.

Downstream analysis and plot generation was performed in R with “Seurat” package. Optimal resolution for clustering was determined using “Clustree” package. Optimal PC for clustering was determined as minimum of where PCs contribute cumulatively to 90% of the standard deviation and where the percent change in variation between the consecutive PCs was less than 0.1% (number of PCs: 16). Previously published gene lists were used to determine signature scores with the “UCell” package.^94^

### FTY720 administration

To prevent lymphocyte egress, mice were injected IP with 1mg/kg FTY720 dissolved in 2% β-hydroxypropyl-cyclodextrin in PBS or vehicle alone. Injections were administered 2 days prior to infection, 5 hours prior to infection (day 0), and subsequently every other day on days 2, 4, and 6 or until lung collection.

### Antigen presenting cell depletion experiments

For depletion of monocytes, 20μg of the anti-CCR2 antibody MC-21 (supplied by Dr. Matthias Mack) or rat IgG2b isotype control were administered IP daily starting 5 hours prior to infection until 5 DPI.

For depletion of alveolar macrophages, mice were treated IN with 100μl of clodronate-loaded or PBS-loaded liposomes daily from -2 DPI to 6 DPI or when lungs were collected. On day of infection, liposome delivery was completed at minimum 5 hours prior to infection.

### MAIT expansion

MAIT cells were expanded with a treatment of CpG (ODN1668, 20μg/mouse) alongside 5-A-RU (50nmol/mouse)+methylglyoxal (1:10 molar ratio of 5-A-RU to methylglyoxal). 5-A-RU+methylglyoxal were incubated for 45 minutes in the dark prior to administration. 5-A- RU+methylglyoxyl was referred to as 5-OP-RU in the text as this is the product of this reaction and previously shown to elicit similar MAIT expansion to synthetic 5-OP-RU.^95^ Lung MAIT expansion was induced by RPh co-administration of CpG and 5-OP-RU on D0, followed by 5-OP-RU only on D1, D2, and D4. Mice were rested for 2 weeks prior to experimentation.

### Co-culture assay

Antigen-presenting cell lines for the co-culture assay included WT3-MR1, DC2.4, DC3.2, and RAW267.4 cells. Murine MAIT hybridoma used was 8D12. All cells were cultured in RPMI 1640 supplemented with 20mM glutaGRO and 10% FBS at 37°C in a 5% CO_2_ incubator. Antigen-presenting cells were seeded at 20,000 cells per 96 well in folate-free RPMI 1640 supplemented with 20mM glutaGRO and 10% FBS four hours prior to addition of bacterial supernatant and 8D12 cells.

For addition of bacterial supernatant to co-culture, overnight cultures of bacteria were backdiluted to an OD of 0.01 and grown at 37°C for three hours. After three hours, an aliquot of culture was used to enumerate CFUs. The remaining culture was centrifuged at 4000rcf for 12 minutes and supernatant was filtered with 0.22μm filter. Bacterial supernatant was added to antigen-presenting cells at 1:100 final dilution after four-hour incubation described above. 8D12 cells were added at 100,000 cells per 96 well. Co-culture was incubated at 37°C in a 5% CO_2_ incubator for 20 hours. Cells were centrifuged at 526rcf for 5 minutes to pellet cells and co-culture supernatant was used for IL-2 ELISA as per manufacturer protocol.

### Fold change for *Mr1^FLAG/+^* mice

Fold change was determined by comparing *Mr1^FLAG/+^* anti-FLAG staining to WT mouse anti-FLAG staining. Both *Mr1^FLAG/+^*and WT mice were subjected to the same infection, digestion protocol, and staining conditions to account for any autofluorescence changes between cell populations and infection model. Median MFI was determined on a biexponential scale. The function for biexponential transformation was extracted utilizing “flowWorkspace” and “CytoML” packages in R. The fold change was determined as biexp(MFI anti-FLAG *Mr1^FLAG/+^*) minus biexp(average anti-FLAG MFI WT) for each cell population. A minimum of three WT staining controls was used for each infection condition.

### Quantification and statistical analysis

Statistical analyses were performed with Prism 10 software (GraphPad). Pairwise comparisons utilized a two-tailed unpaired Student’s *t* test with Holm-Šidák multiple comparison correction if data were normally distributed following a Shapiro-Wilk normality test or a Mann-Whitney test with Holm-Šidák multiple comparison correction if data were not normally distributed. Normality to determine appropriate test was determined using Kruskal-Wallis test. Analyses with more than 2 groups utilized a one-way ANOVA with Šidák multiple comparison correction. Survival statistics were performed using a Mantel-Cox test. * = *P* < 0.05, ** = *P* < 0.01, *** = *P* < 0.001, and **** = *P* < 0.0001. “ns” = not significant.

## STAR Methods

### Key Resources Table

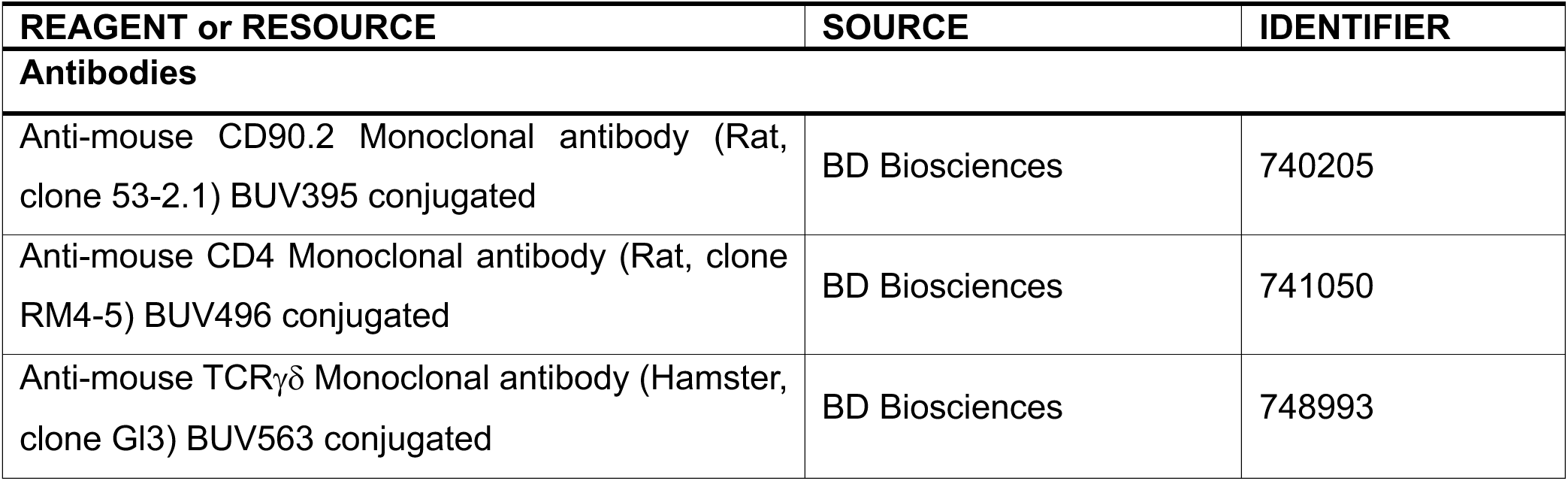
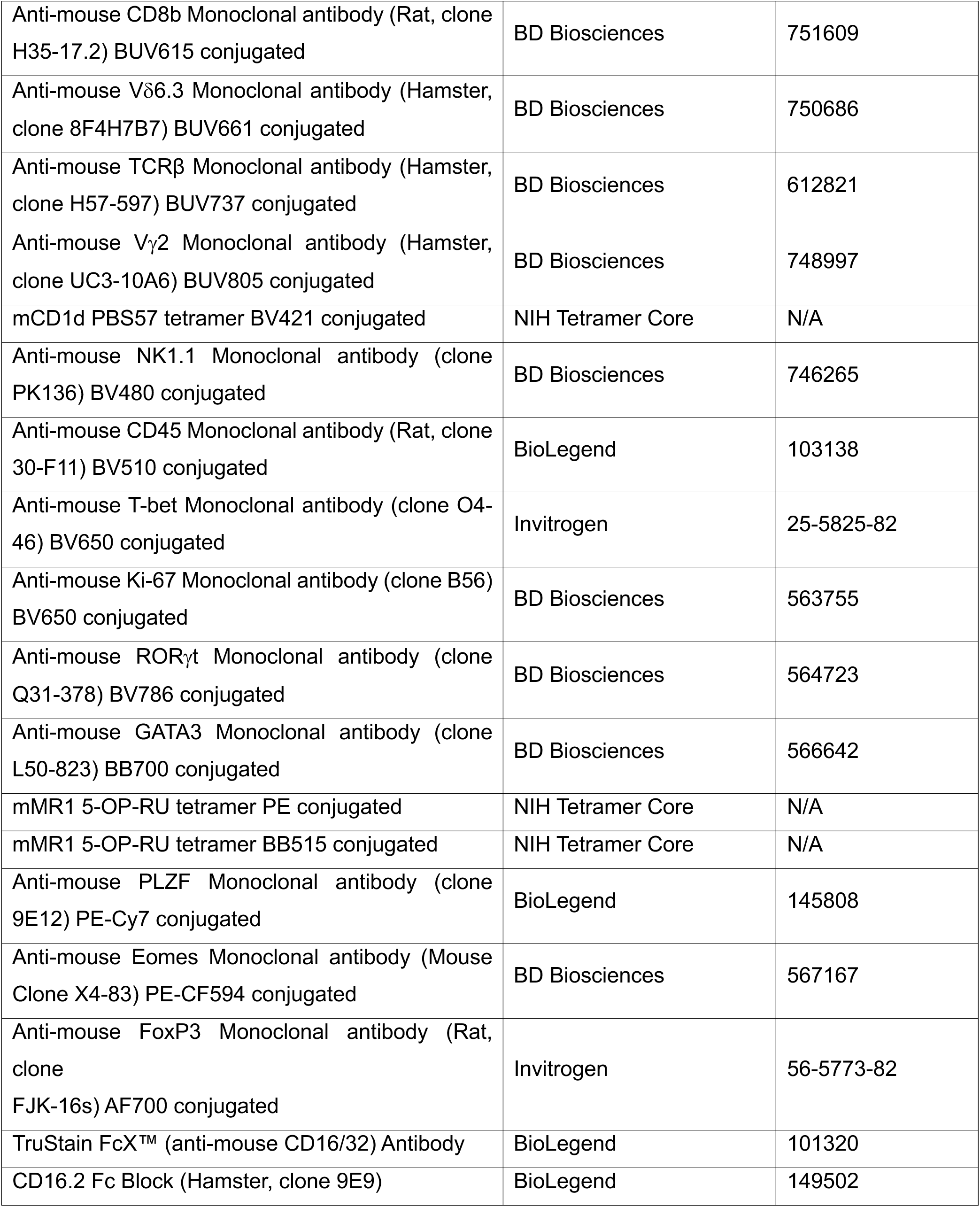
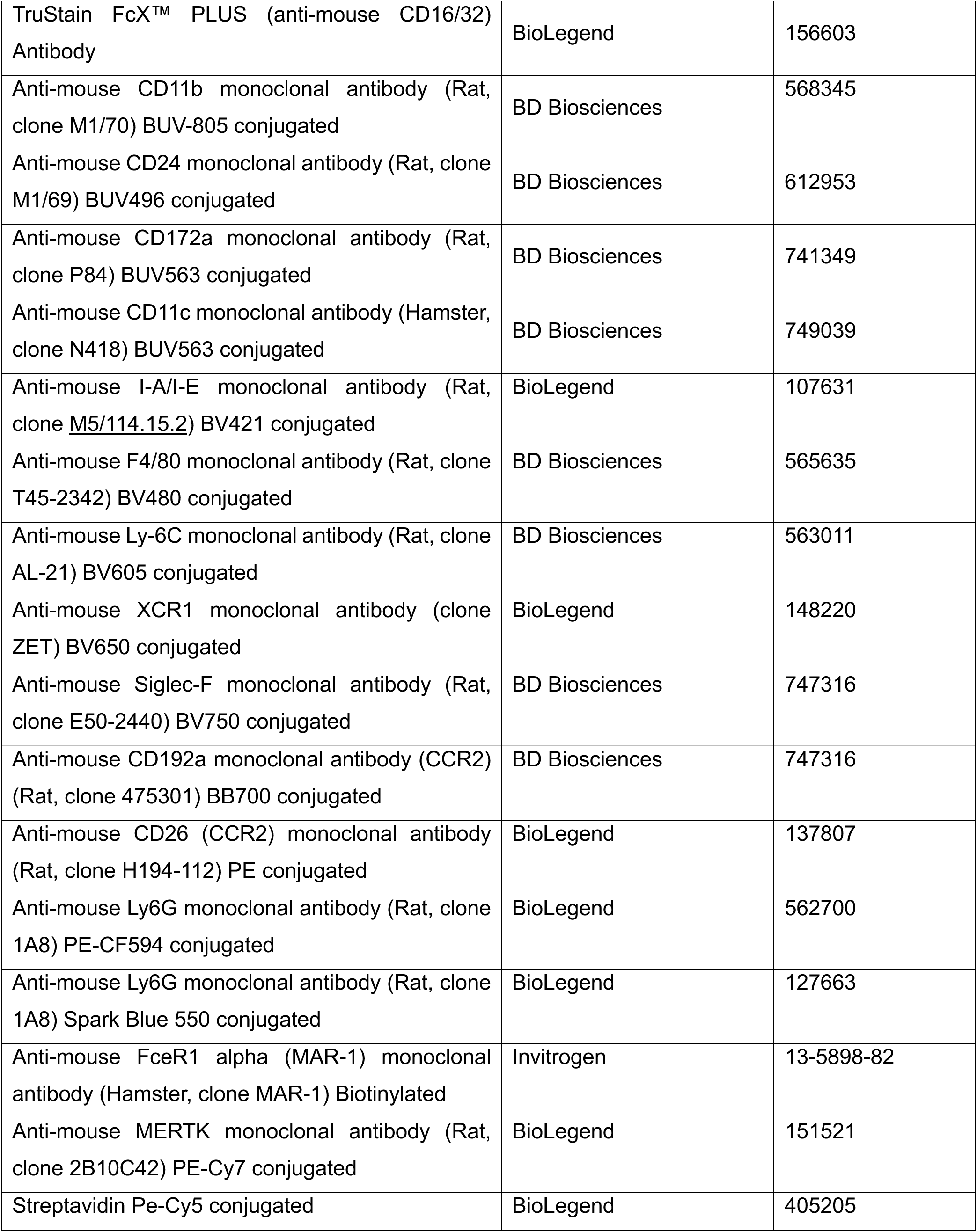
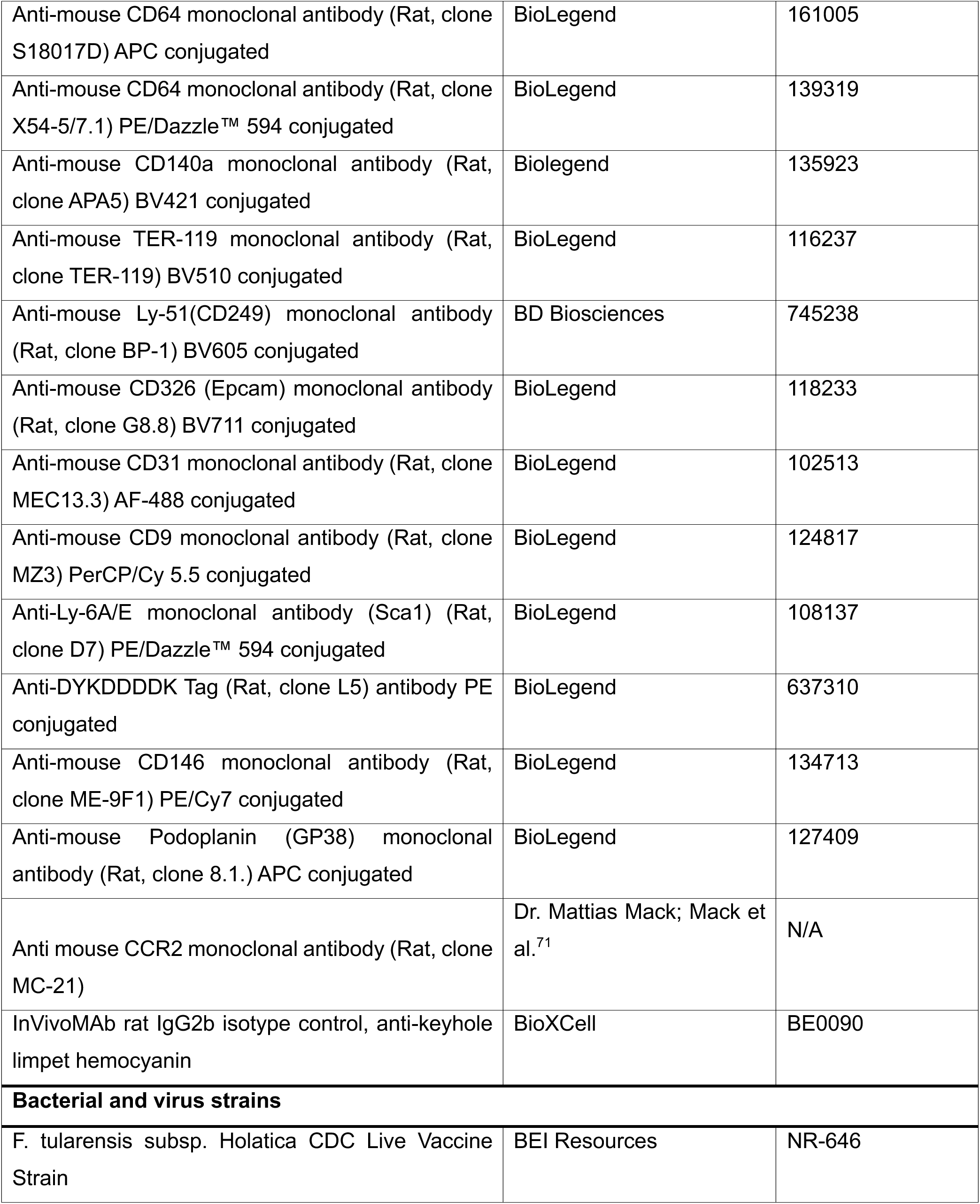
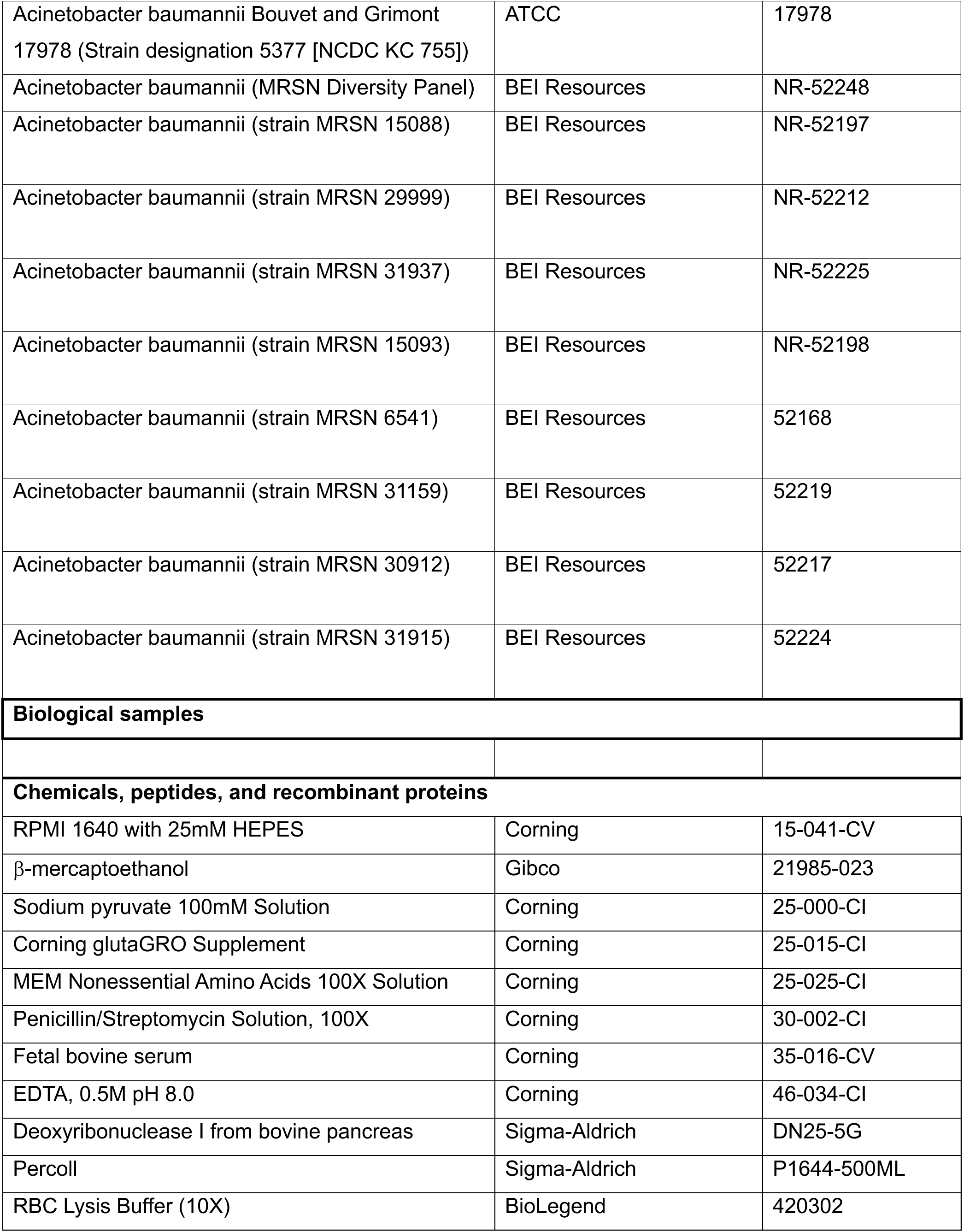
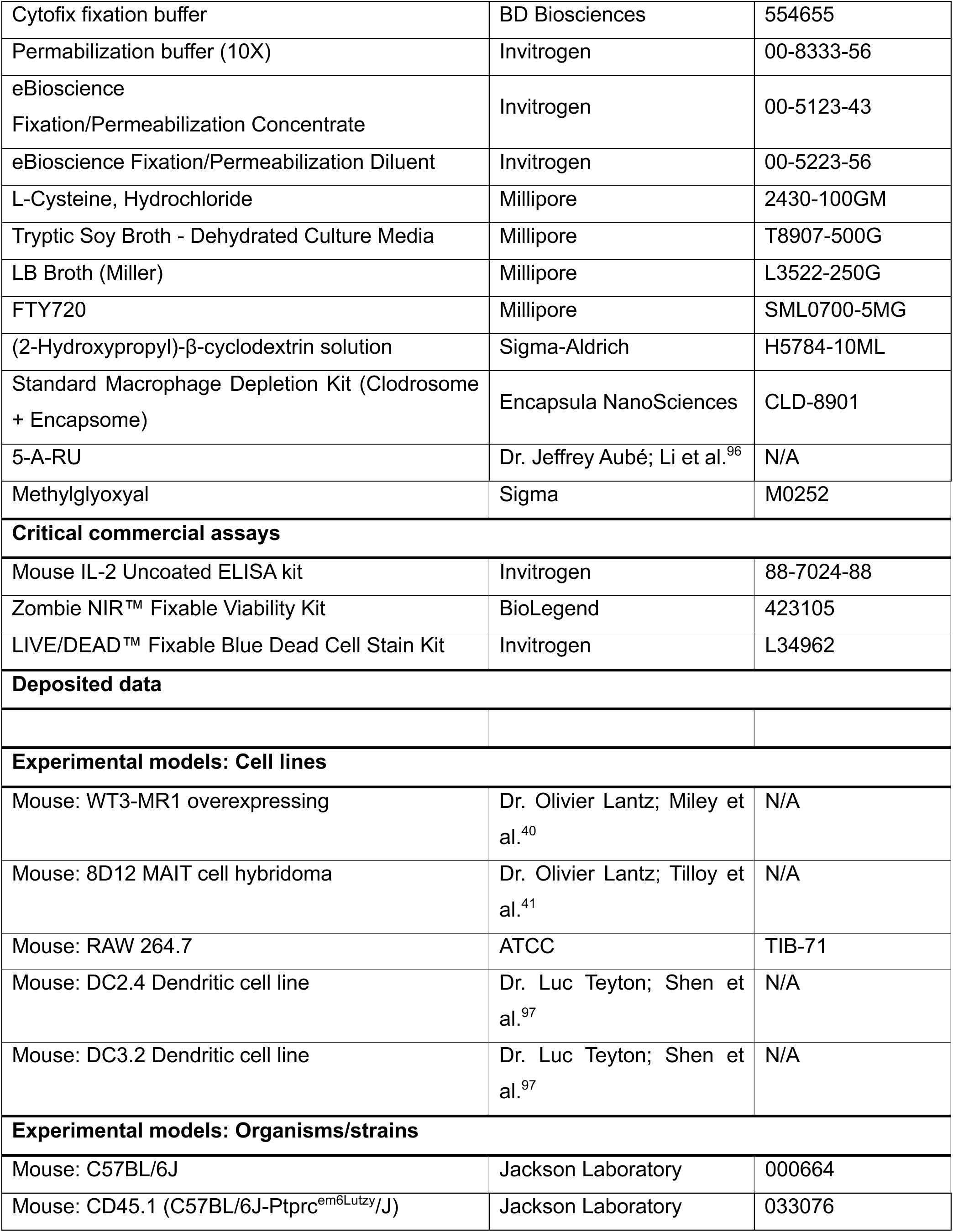
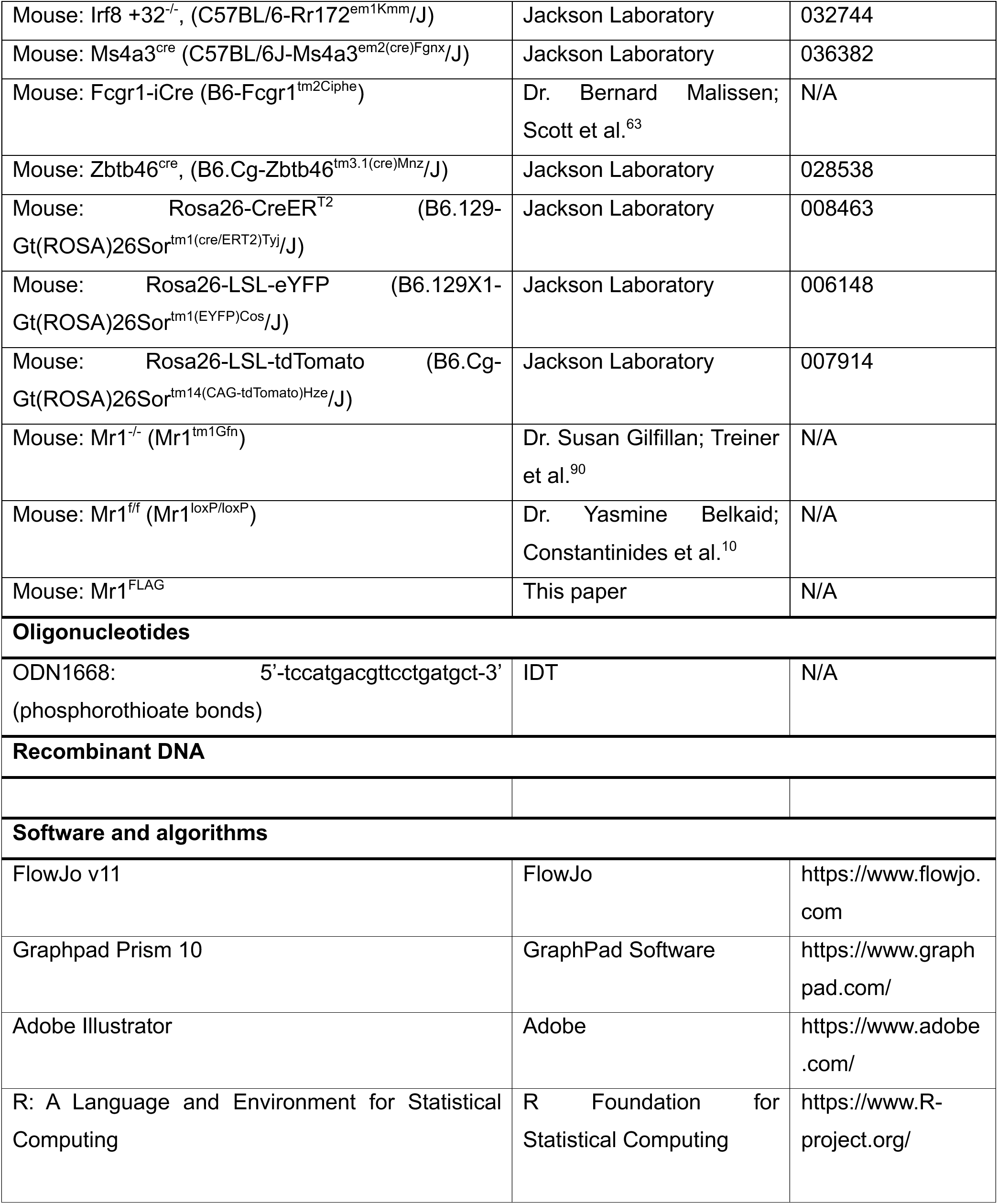
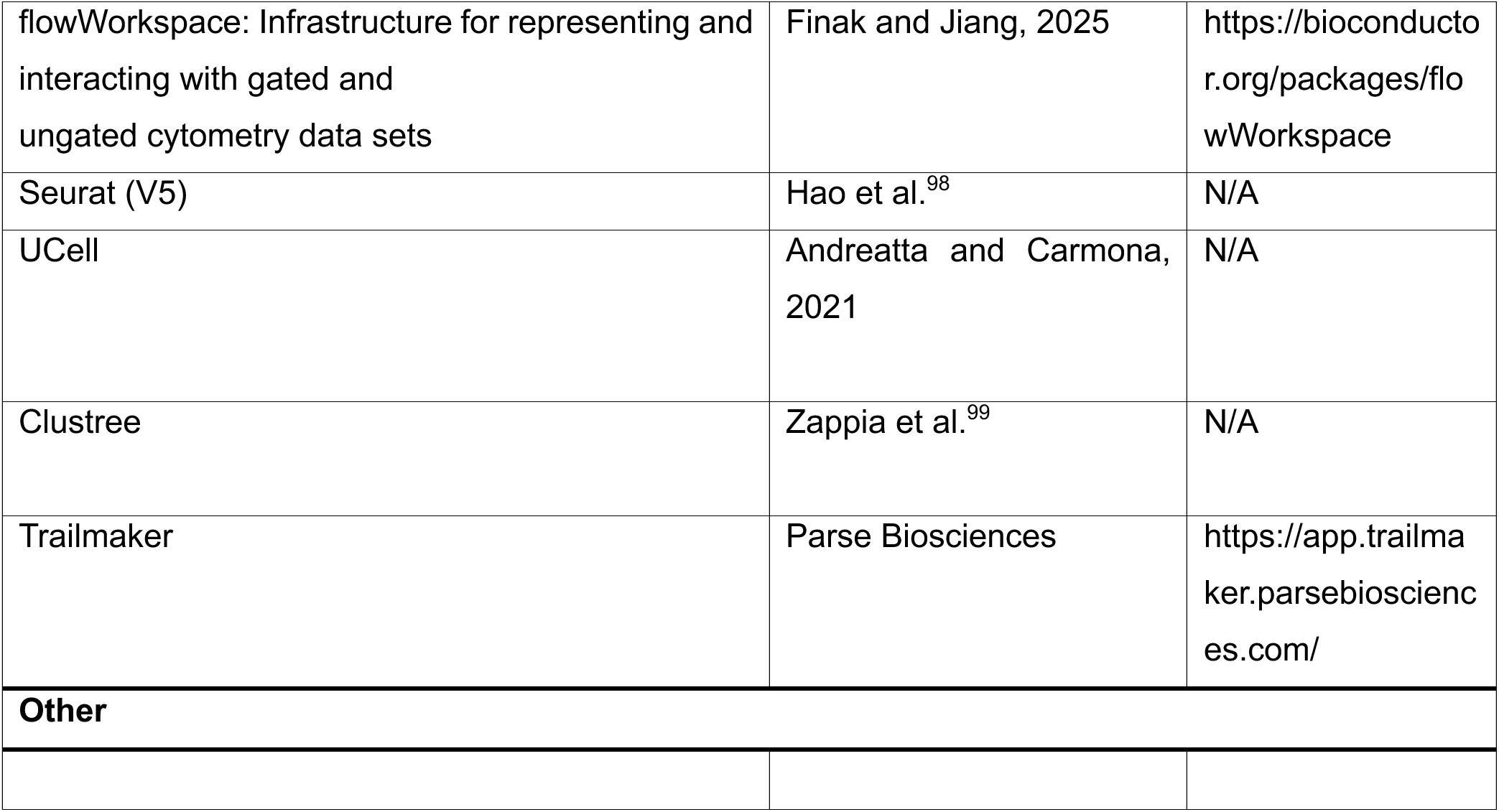

---

**Figure S1.**
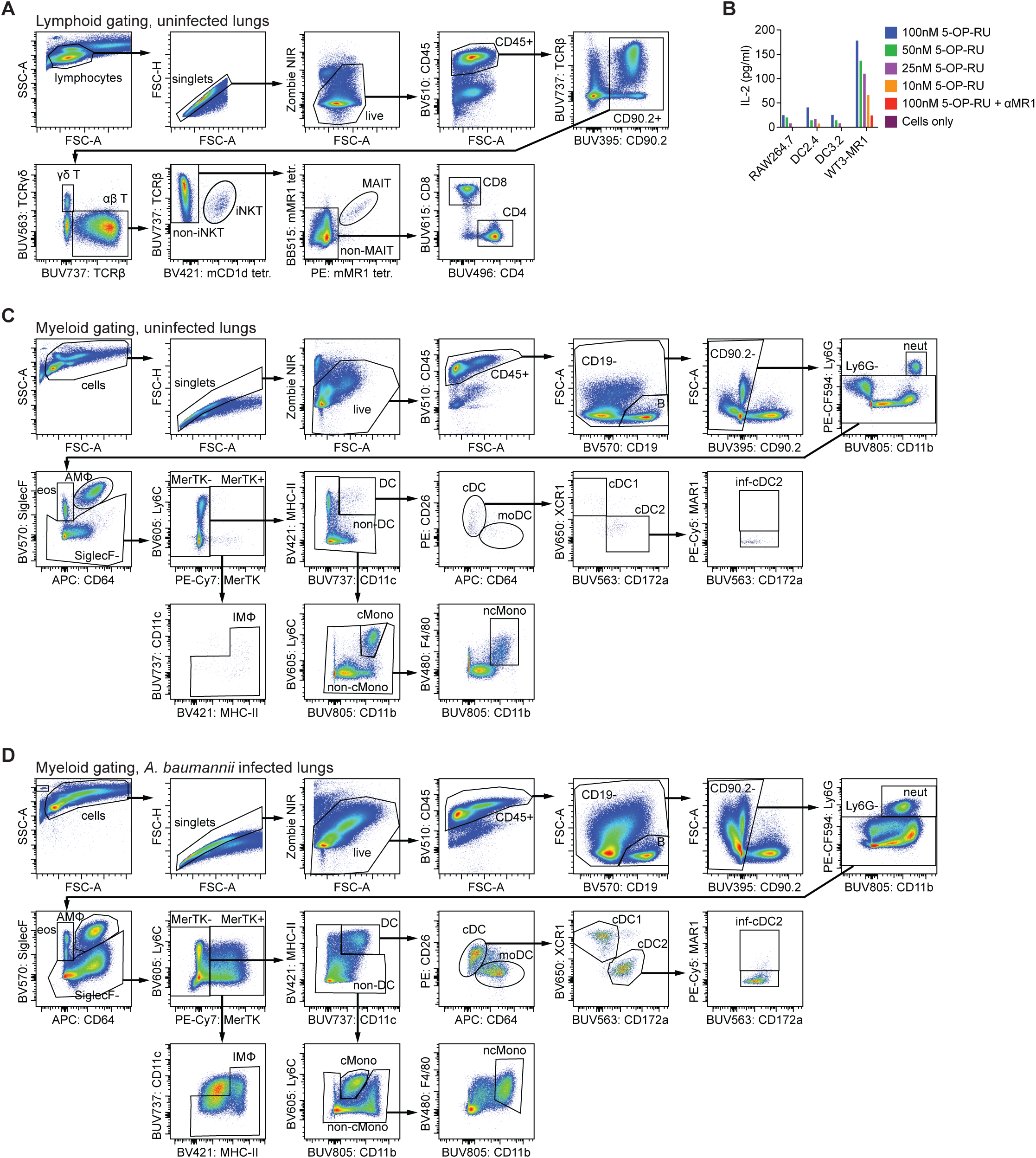
Flow cytometry gating and co-culture validation. (A) Lymphoid gating scheme for lungs. Gates for lymphocytes remain the same in infection conditions. (B) Titration of 5-OP-RU in 8D12 and antigen-presenting cell co-culture across different antigen-presenting cell lines. (C) Myeloid gating scheme for uninfected lungs. Ly6C^+^ cMonos were additionally gated CCR2^+^ and minor adjustments were made to accommodate the PE-FLAG antibody, when necessary. (D) Myeloid gating scheme for *A. baumannii* infected lungs. Ly6C^+^ cMonos were additionally gated CCR2^+^ and minor adjustments were made to accommodate the PE-FLAG antibody, when necessary.

**Figure S2.**
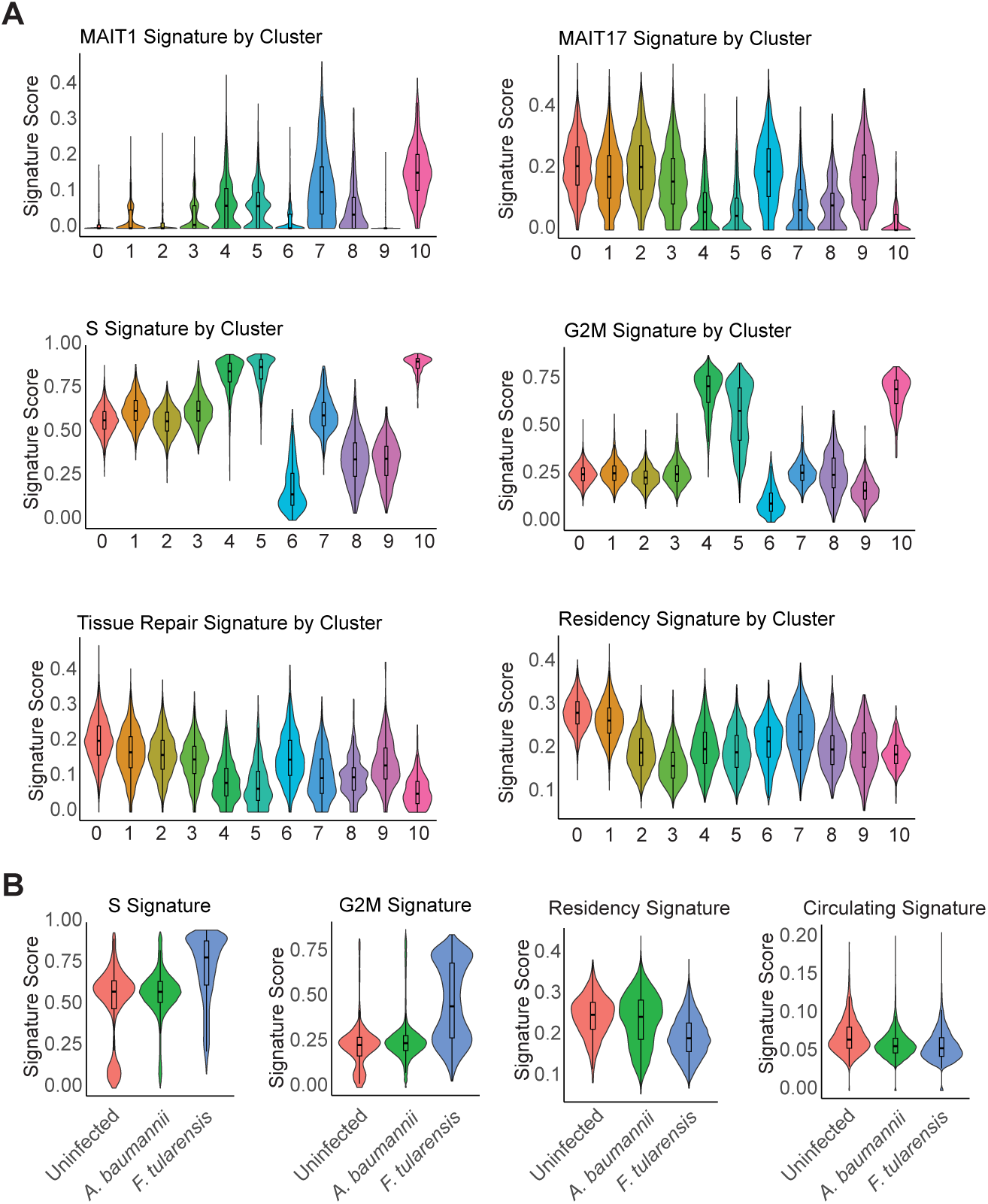
Transcriptional profiling of pulmonary MAIT cells. scRNA-seq data of MAIT cells sorted from uninfected (1482 cells), 7 DPI RPh *A. baumannii* (11690 cells), and 7 DPI RPh *F. tularensis* (5470 cells) using 10X Genomics platform. Plots were generated by combining scRNA-seq libraries from these three samples. (A) Enrichment for MAIT1 signature, MAIT17 signature, S phase signature, G2M phase signature, tissue repair signature, and tissue residency signature split by cluster. (B) Enrichment for S phase signature, G2M phase signature, tissue residency signature, and circulating signature split by sample.

**Figure S3.**
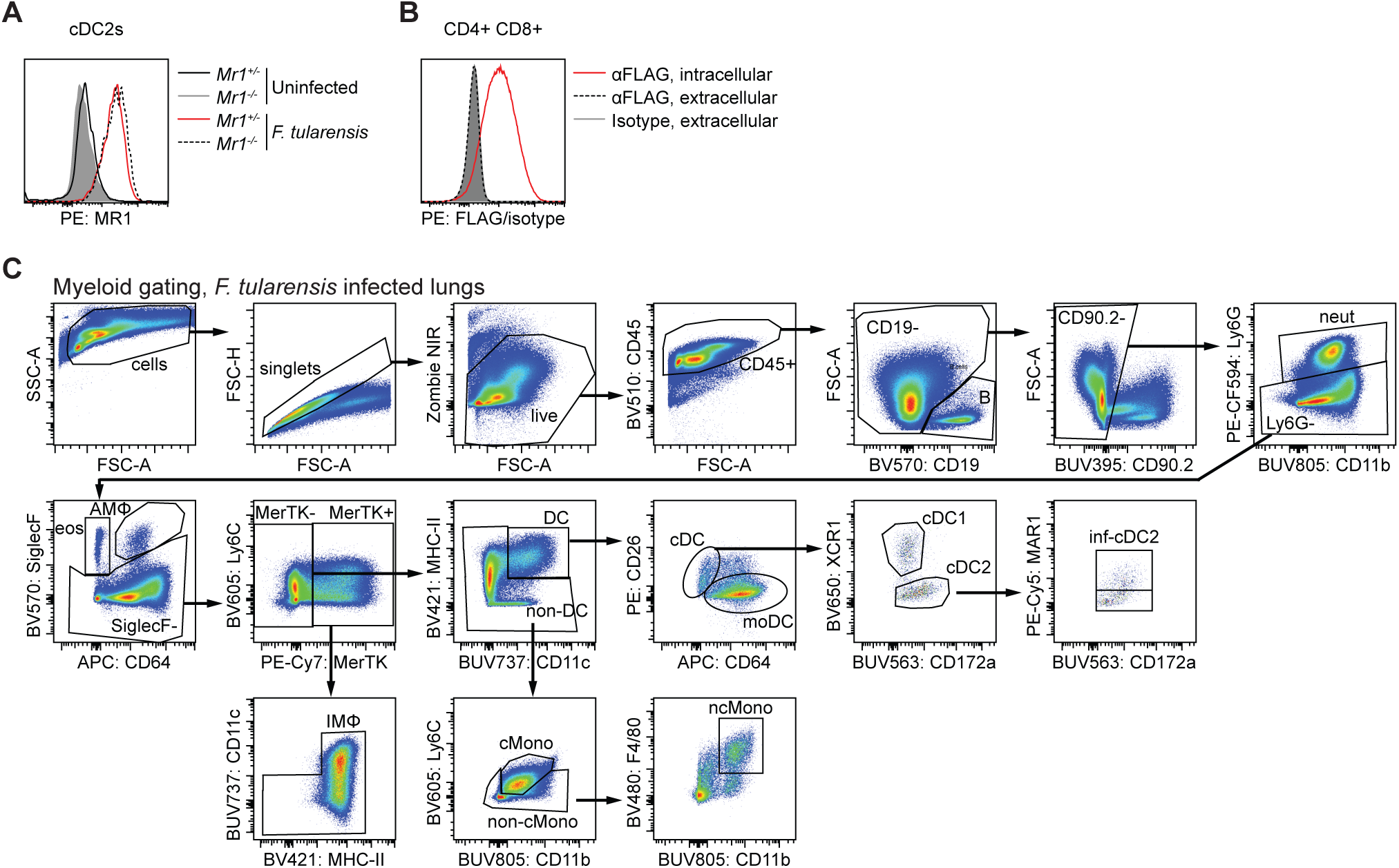
Validation of *Mr1^FLAG^* strain. (A) Representative flow staining for extracellular αMR1 (26.5 clone) on lung cDC2s between uninfected and 6 DPI *F. tularensis* for both *Mr1*^+/-^ and *Mr1*^-/-^ mice. (B) Representative flow staining for αFLAG and isotype on CD4^+^ CD8^+^ thymocytes compared between extracellular and intracellular stain. (C) Myeloid gating scheme for *F. tularensis* infected lungs. Ly6C^+^ cMonos were additionally gated CCR2^+^ and minor adjustments were made to accommodate the PE-FLAG antibody, when necessary.

**Figure S4.**
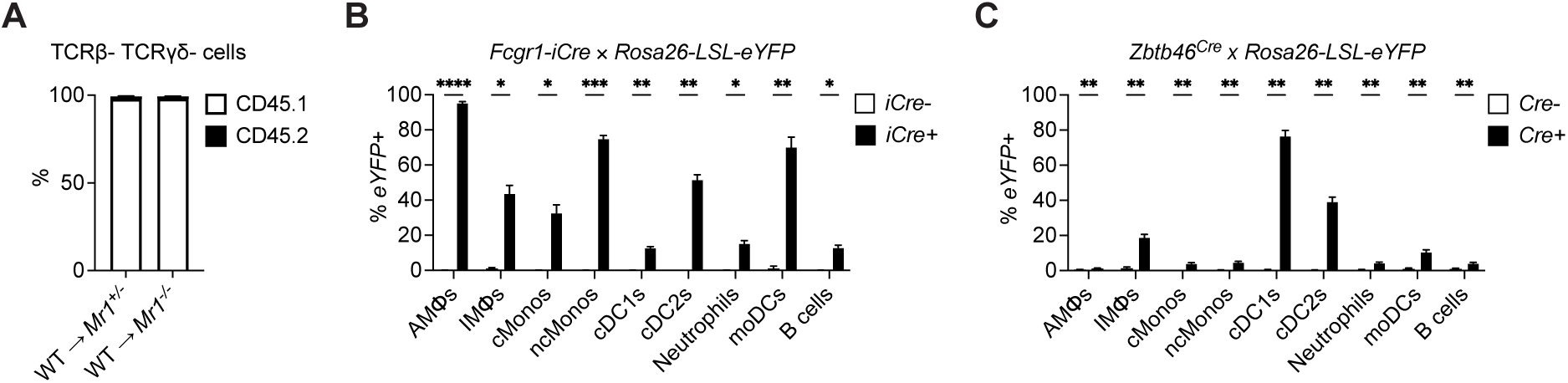
Verification of bone marrow reconstitution and expression of *Fcgr1-iCre* and *Zbtb46^Cre^* alleles. (A) Reconstituted and residual live, TCRβ^-^, TCRγδ^-^ cells identified using congenic markers (CD45.1 and CD45.2) in the lungs of WT *→ Mr1^+/-^* and WT *→ Mr1^-/-^* chimeras (n=12-13/group). (B) YFP expression in *Mr1^f/f^ Fcgr1^iCre^* × *Rosa26-LSL-eYFP* compared to *Mr1^f/f^*× *Rosa26-LSL- eYFP* littermate controls (n=4-5/group). (C) YFP expression in *Mr1^f/f^ Zbtb46^Cre^* × *Rosa26-LSL-eYFP* compared to *MR1^f/f^*× *Rosa26-LSL- eYFP* littermate controls (n=7-8/group) Data is representative of two or more independent experiments. Graphs indicate means ±SEM. Unpaired Student’s *t*-test (B-C) was performed. Statistics: ns, not significant (*P* > 0.05); \**P* < 0.05; \*\**P* < 0.01; \*\*\**P* < 0.001, \*\*\*\**P* < 0.0001

**Figure S5.**
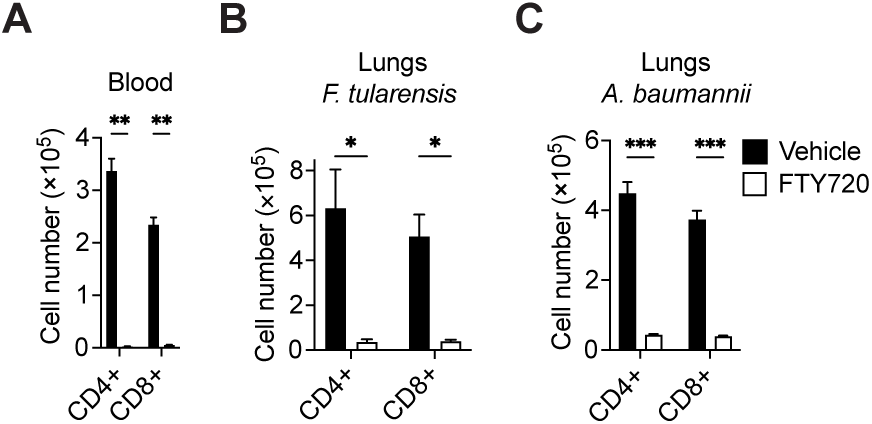
Effect of FTY720 treatment on conventional T cells. (A) Number of CD4^+^ and CD8^+^ T cells in blood 1 day after treatment of vehicle or FTY720 (n=3/group). (B) Number of CD4^+^ and CD8^+^ T cells in the lungs of vehicle or FTY720 treated mice either uninfected or 7 DPI *F. tularensis* (n=5/group). (C) Number of CD4^+^ and CD8^+^ T cells in the lungs of vehicle or FTY720 treated mice either uninfected or 7 DPI *A. baumannii* (n=5-6/group). Data is representative of two or more independent experiments. Unpaired Student’s *t*-test (A-C) was performed. Statistics: ns, not significant (*P* > 0.05); \**P* < 0.05; \*\**P* < 0.01; \*\*\**P* < 0.001, \*\*\*\**P* < 0.0001

**Figure S6.**
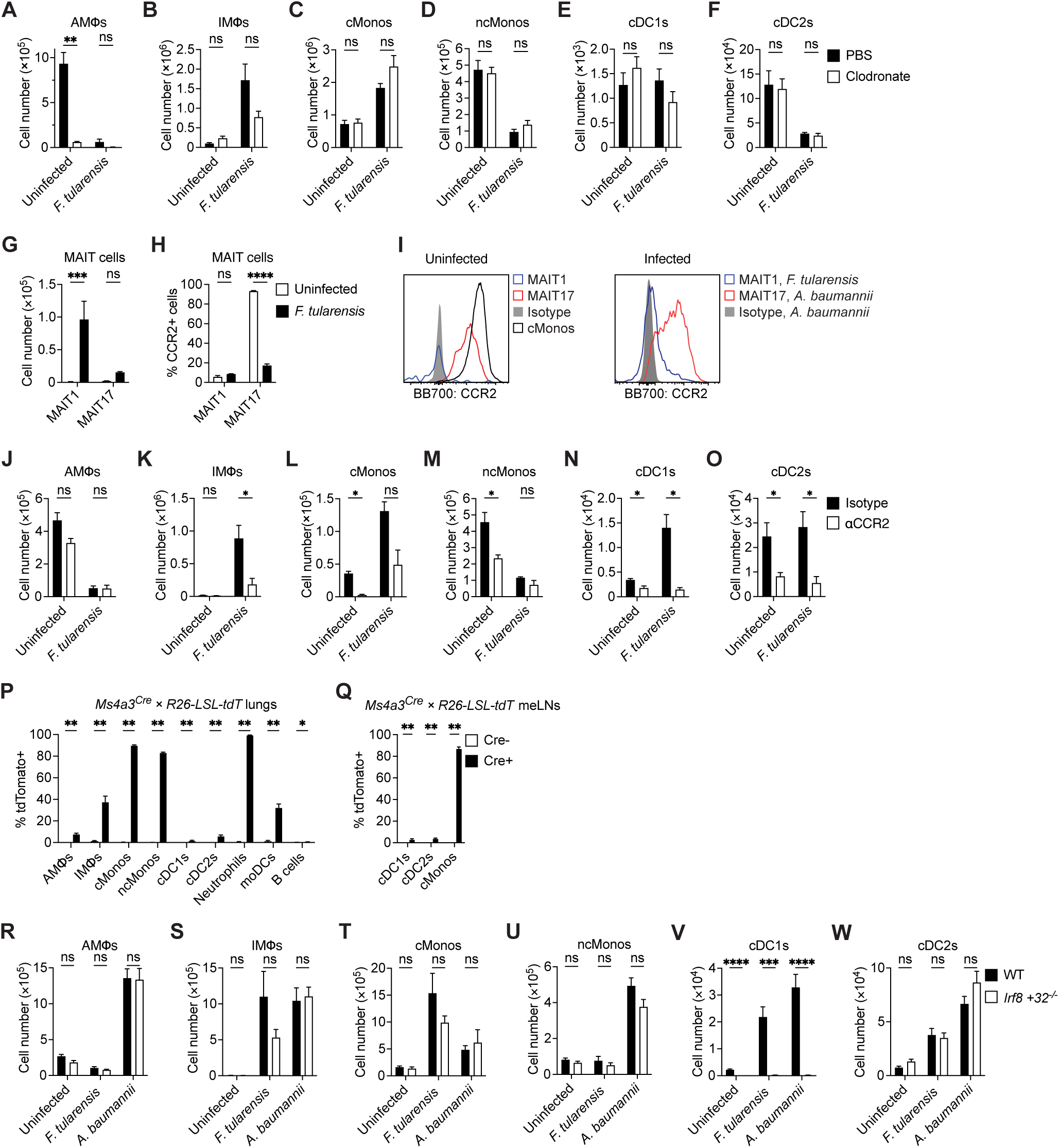
Confirmation of myeloid cell depletion and *Ms4a3^Cre^* expression. (A-F) Number of AMΦs (A), IMΦs (B), cMonos (C), ncMonos (D), cDC1s (E) and cDC2s (F) from the lungs of PBS or clodronate treated mice either uninfected or 7 DPI *F. tularensis* (n=4-6/group). (G-H) Number of MAIT1 and MAIT17 cells (G) and their respective CCR2 expression (H) in mice uninfected or 7 DPI *F. tularensis* (n=6/group). (I) Representative histogram displaying CCR2 expression of cMonos (black), MAIT17 (red), MAIT1 (blue) and bulk MAIT cells stained with isotype control (filled grey) analyzed in uninfected or infected mice as indicated. (J-O) Number of AMΦs (J), IMΦs (K), cMonos (L), ncMonos (M), cDC1s (N) and cDC2s (O) from the lungs of mice treated with IP αCCR2 or isotype control either uninfected or 7 DPI *F. tularensis* (n=5/group). (P) tdTomato expression in the lungs of *Ms4a3^Cre^ Rosa26-LSL-tdTomato* compared to *Rosa26-LSL-tdTomato* controls (n=6-7/group). (Q) tdTomato expression in the meLNs of *Ms4a3^Cre^ Rosa26-LSL-tdTomato* compared to *Rosa26-LSL-tdTomato* controls (n=6-7/group). (R-W) Number of AMΦs (R), IMΦs (S), cMonos (T), ncMonos (U), cDC1s (V) and cDC2s (W) in the lungs of WT and *Irf8 +32^-/-^* mice either uninfected or 7 DPI *F. tularensis* or 7 DPI *A. baumannii.* (n=4-11/group). Data is representative of two or more independent experiments. Graphs indicate means ±SEM. Unpaired Student’s *t*-test (A, B, D, E, H, J, M, R, T, U) or Mann-Whitney U test (C, F, G, K, L, N, O, P, Q, S, V, W) was performed. Statistics: ns, not significant (*P* > 0.05); \**P* < 0.05; \*\**P* < 0.01; \*\*\**P* < 0.001, \*\*\*\**P* < 0.0001.

